# Endophyte inoculation enhances *Ulmus minor* resistance to Dutch elm disease

**DOI:** 10.1101/2020.05.04.076349

**Authors:** C Martínez-Arias, J Sobrino-Plata, S Ormeño-Moncalvillo, L Gil, J Rodríguez-Calcerrada, JA Martín

## Abstract

Certain fungal endophytes are known to improve plant resistance to biotic stresses in forest trees. In this study, three stem fungal endophytes belonging to classes Cystobasidiomycetes, Eurotiomycetes and Dothideomycetes were selected from 210 isolates for their potential as enhancers of *Ulmus minor* resistance to *Ophiostoma novo-ulmi*. We evaluated phenotypic traits of these endophytes that could be beneficial for inhibiting *O. novo-ulmi* in the host plant. Under *in vitro* conditions, the Dothideomycetous isolate YCB36 strongly inhibited *O. novo-ulmi* growth, released antipathogenic VOCs, chitinases and siderophores, and overlapped with the pathogen in nutrient utilization patterns. These functional traits could explain the 40% reduction in leaf wilting due to *O. novo-ulmi* in elm trees pre-inoculated with this endophyte. *Ulmus minor* trees inoculated with this endophyte showed increased leaf stomatal conductance and higher concentrations of flavonoids and total phenolic compounds in xylem tissues, suggesting induction of defence metabolism.

## 1. Introduction

In natural ecosystems, plants host a broad spectrum of microbes, especially fungi and bacteria that can colonize the plant surface as epiphytes or inner plant tissues as endophytes. When living inside plants, some endophytes act as mutualistic symbionts, assisting plant functioning, survival and fitness (Carroll, 1988; Schulz and Boyle, 2005; Rodríguez et al., 2009; Hardoim et al., 2015; Vandenkoornhuyse et al., 2015). Fungal endophytes have received considerable attention in recent years because of their impact on plant fitness and plant responses to biotic and abiotic stressors (Singh et al., 2011; Lata et al., 2018; Rabiey et al., 2019), and for being a potential source of bioactive compounds (Richardson et al., 2015; McMullin et al., 2018; Tanney et al., 2018). The role of fungal endophytes in plant resilience is especially important in trees, which face extreme weather conditions and recurring pathogen and insect attacks during their long lifespan (Lau et al., 2017; Terhonen et al., 2019, Martínez-Arias et al., 2020).

Endophytic fungi can shape host resistance to biotic stressors through several mechanisms (Witzell and Martín, 2018). Some of these are based on direct interaction between endophytes and pathogens, such as mycoparasitism, competitive exclusion by substrate consumption, and inhibition of the pathogen by extracellular enzymes released by the endophyte, e.g. by proteases or chitinases, which are able to degrade the hyphal cell wall of pathogens (Guthrie and Castle, 2006; dos Reis Almeida et al., 2007; Orlandelli et al., 2015). Indirectly, endophytes can stimulate the defence response of the plant by altering different signal transduction pathways, sometimes related with jasmonic or salicylic acid signalling(Shoresh et al., 2010; Fesel and Zuccaro, 2016; Martínez-Medina et al., 2017; Khare et al., 2018; Naidoo et al., 2019). Some endophytes enhance plant growth by synthesizing gibberellins (e.g. gibberellic acid, GA_3_) and auxins (e.g. indole-3-acetic acid, IAA) (Patten and Glick, 2002; Shi et al., 2009; Doty et al., 2011; Videira et al., 2012) or by improving nutrient acquisition (Della Monica et al., 2015; Surono and Narisawa, 2017; He et al., 2019), helping to counteract the negative effects of pathogen invasion. Phosphate solubilization (Nutaratat, 2014; Priyadharsini et al., 2017), siderophore emission (Moller et al., 2016; Haselwandter et al., 2020) and hydrogen cyanide (HCN) production (Haas and Defago, 2005) are the main mechanisms by which symbionts regulate availability of nutrients in soil. Siderophores and HCN have also been reported to aid biological control of plant pathogens by acting as antimicrobial compounds (Haselwandter et al., 2007; Rijavec and Lapanje, 2016). Endophyte-induced oxidative stress could enhance synthesis of plant antioxidant molecules and increase oxidative protection from biotic and abiotic stressors (White and Torres, 2010). Flavonoids and other phenolic antioxidants are some of the main molecules involved in these processes (Yang et al., 2016).

In recent decades, interest in the study of fungal endophytes and their use as biological control agents has increased (Arnold et al., 2003; Mejía et al., 2008; Fakhro et al., 2010; Martínez-Álvarez et al., 2016; Martínez-Arias et al., 2019; Quiring et al., 2019; Rabiey et al., 2019; Halecker et al., 2020). In forest trees, a successful example of biological control with fungal endophytes is the case of *Phialocephala scopiformis*, a rugulosin-producing endophyte with the ability to reduce the incidence of the *Picea glauca* budworm (Miller et al., 2008; Quiring et al., 2019). Effective applications of biocontrol agents require, preferably, locally adapted endophytes and easily cultured strains. Using local endophytes reduces the risk of incompatible interactions with the host plant and adverse environmental conditions for fungal development as well as higher and long-term fungal stability (Frasz et al., 2014); while easily cultured endophytes facilitate inoculum production. Multi-species microbial consortia can provide better results than single species due to the wider array of complex functions they perform (Brenner et al., 2008). However, studying a single species or a consortium of a few species is a necessary first step to evaluate specific functions and effects.

Dutch elm disease (DED), initially caused by *Ophiostoma ulmi*, was introduced into Europe and North America by global trade in the first half of the 20th century, being one of the earliest examples of alien diseases threatening native forests. The current DED pandemic is caused by *Ophiostoma novo-ulmi* (Sordariomycetes), which is transmitted into healthy elm trees through bark beetles of the genera *Scolytus* and *Hylurgopinus*. *O. novo-ulmi* spreads within xylem vessels, causing blockage and embolism (Brasier and Buck, 2001; Martín et al., 2019). Among the various biocontrol agents studied to reduce the impact of DED (Scheffer et al., 1990; Solla and Gil, 2003), fungal endophytes have received increasing attention in recent years (Martín et al., 2013; Blumenstein et al., 2015; Martín et al., 2015). In elms, distinct fungal assemblages were found in the xylem of susceptible and tolerant genotypes (Martín et al., 2013). Some of these endophytes were isolated and tested against *O. novo-ulmi in vitro* and in field plot assays (Martín et al., 2015), and some showed the ability to reduce DED symptoms. However, environmental factors and host genotype appeared to alter the behaviour and spread of the inoculated fungi, providing an unstable protective effect (Martín et al, 2015).

In this work, we aimed to evaluate three additional elm fungal endophytes for their use as biocontrol candidates against DED. For this purpose, we analyzed parameters that could be relevant for the successful inhibition of the pathogen by endophytes or for increasing host resistance, in both cases decreasing host damage. We selected three fungal endophytes from a collection of 210 isolates following two criteria. The first criterion was based on earlier metabarcoding research by our group that revealed a close association between the abundance of operational taxonomic units within classes Cystobasidiomycetes and Eurotiomycetes and the level of host resistance to DED (Macaya-Sanz et al., 2020). Considering this result, we hypothesize that elm endophytes belonging to these fungal classes exhibit functional traits involved in *U. minor* resistance to DED. The second criterion was based on other research showing that although most endophytes are able to reduce the growth of *O. novo-ulmi* in *in vitro* dual tests to some extent, mainly through substrate competition or weak antibiosis, only a few endophytes are able to strongly inhibit pathogen growth by antibiosis (Martín et al., 2015). Therefore, we further hypothesize that fungal endophytes exhibiting strong antibiotic activity towards *O. novo-ulmi* have the potential to reduce *O. novo-ulmi* spread *in planta* and consequently decrease disease symptoms.

Two endophytes representative of classes Cystobasidiomycetes and Eurotiomycetes, and an additional Dothideomycetous isolate which showed strong *in vitro* antibiosis towards *O. novo-ulmi* were selected for this work. To evaluate the biocontrol potential of these endophytes we performed four specific studies to: i) characterize *in vitro* the type of interaction and the antibiotic activity of the three selected endophytes against *O. novo-ulmi*, ii) determine the nutrient metabolism, nutrient uptake ability and production of plant growth-promoting compounds *in vitro* by the endophytes, iii) evaluate the plant colonization ability of the endophytes and their protective effect against *O. novo-ulmi* in a field assay with 6-year-old susceptible *U. minor* trees, and iv) study the plant functional and chemical response to the endophytes to explain the protective effect against the pathogen.

## 2. Material and methods

### 2.1 Fungal material

Three fungal endophytes isolated from elm twigs and named YM11, P5 and YCB36 were selected for the experiments. The strains were deposited in the Spanish Type Culture Collection (CECT) with the codes CECT 13193, CECT 13192, and CECT 21178, respectively. The endophytes are from a collection of fungi that includes 210 isolates obtained from *U. minor* trees in the last 10 years, conserved by the Spanish elm breeding programme. The three selected fungi were isolated in 2014 and 2015 from three *U. minor* genotypes tolerant to DED (Martín et al., 2015) growing in a conservation plot at “Puerta de Hierro” Forest Breeding Centre, Madrid (Spain, 40° 27’ 24’’ N; 3° 45’ 0’’ W; 600 m. a.s.l.). Twig samples for fungal isolation were first deeply cleaned with sterilized distilled water, and then sterilized following the method III described in Schulz et al. (1993). The yeasts belong to the Eurotiomycetes (YM11) and the Cystobasidiomycetes (P5), two fungal classes commonly occurring in *U. minor* trees with high resistance to *O. novo-ulmi* (Macaya-Sanz et al., 2020). YM11 colonies growing in standard culture medium comprise a mixture of budding cells and hyphae, while P5 forms bud cell aggregates. Additionally, the YCB36 isolate, a filamentous fungus from the Dothideomycetes, was selected because it strongly inhibited *O. novo-ulmi* growth in preliminary *in vitro* dual culture screening tests (visual assessment). This fungus produces a dark halo around the colony when grown on 3.5 % yeast extract agar (YEA), indicating release of metabolites that could cause inhibition in *O. novo-ulmi* growth. YCB36 had an *in vitro* growth of 3.89 mm per day when growing in YEA.

After initial isolation on YEA, 4 × 4 mm plugs of each endophyte isolate were conserved in sterilized distilled water (dH_2_O) at 4 ° C in the dark. Two months before the experiment, isolates were sub-cultured on YEA medium in the dark at 22 ° C. These colonies were subcultured every 15 days onto fresh YEA medium and used as stock source for subsequent assays. For molecular identification of endophytes, isolates were grown on YEA medium over an autoclaved cellophane layer. Five hundred milligrams of the mycelium were harvested and ground with a pellet pestle. DNA extraction was carried out as described by Martínez-Arias et al. (2019). After DNA extraction, the two Internal Transcribed Spacers (ITS) of ribosomal DNA and the large subunit of the rRNA (LSU) were amplified through Polymerase Chain Reaction (PCR) for each fungal strain. The universal primers ITS1F (5’-CTTGGTCATTTAGAGGAAGTAA-3’) and ITS4 (5’- TCCTCCGCTTATTGATATGC-3’) were used to amplify the ITS region following a protocol based on White and Bruns (1990). In the case of the LSU region, the primers LROR (5’- ACCCGCTGAACTTAAGC-3’) and LR5 (5’-ATCCTGAGGGAAACTTC-3’) were selected according to Vilgalys and Hester (1990). In addition to the previous markers, some protein-coding genes were used for YM11 and YCB36 identification. Thus, the translation elongation factor 1-alpha (*TEF1α*) was amplified through the primers EF1-1018F (5’- GAYTTCATCAAGAACATGAT-3’) and EF1-1620R (5’ GACGTTGAADCCRACRTTGTC-3’) (Stielow et al., 2015). YCB36 taxonomical assignment was also characterized through amplification of beta-tubulin (*tub2/BenA*) with the primers Bt2a (5’- GGTAACCAAATCGGTGCTGCTTTC-3’) and Bt2b (5’- ACCCTCAGTGTAGTGACCCTTGGC-3’) (Glass and Donaldson, 1995), and the amplification of the gene for partial actin with the primers act- 512F (5’- ATGTGCAAGGCCGGTTTCGC-3’) and act-738R (5’- TACGAGTCCTTCTGGCCCAT-3’) (Carbone and Kohn, 1999). The amplification products were Sanger-sequenced, and sequences were annotated taxonomically based on the most similar sequence with taxonomic identity using the Genbank Basic Local Alignment Search Tool (BLASTn) of the National Centre for Biotechnology Information database (NCBI, MD, USA).

The *O. novo-ulmi* ssp. *americana* isolate SOM-1 used in the experiments (Martín et al., 2019) was isolated in 2014 from a 168-year-old *U. minor* tree in El Pardo (Madrid, Spain) and had an *in vitro* growth rate of 4 mm per day on 2% malt extract agar (MEA) at 20 ° C. The identity of the pathogen was determined by assessments of colony morphology and growth rates at 20 ° C and 33 ° C, according to Brasier (1981). To confirm species and subspecies identity, a molecular analysis of the isolate was performed according to Gibb and Hausner (2005), as described in Martín et al. (2019). The isolate was conserved as mycelial plugs immersed in sterilized dH_2_O and subcultured on MEA two months before the experiment, kept in the dark at 22 ° C, and subcultured every 15 days.

### 2.2 In vitro characterization of fungal endophytes

#### 2.2.1 Dual culture with O. novo-ulmi

Each endophyte was co-cultured with *O. novo-ulmi* in petri dishes (90 mm in diameter). Mycelial plugs (5 x 5 mm) of both the endophyte and the pathogen were subcultured from the actively growing colony edge and transferred to new dishes with YEA. *Ophiostoma novo-ulmi* and endophyte plugs were spaced 4 cm apart in the petri dish. In control plates, *O. novo-ulmi* was grown as a monoculture on one side of the petri dish. Three replicate plates per endophyte (dual cultures) and pathogen (monocultures) were used. Colony growth was measured at 3, 7, 10 and 12 days of incubation by scanning the growth area of each fungus using ImageJ software (http://imagej.nih.gob/ij/). Relative growth inhibition of *O. novo-ulmi* by endophytes was calculated by comparing *O. novo-ulmi* colony growth in control plates and endophyte- confronted plates (Chamberlain and Crawford, 1999; Kusari et al., 2013). The following interactions between the endophyte and the pathogen were also assessed qualitatively according to Mejía et al. (2008): i) antibiosis (the endophyte is able to reduce pathogen growth by the release of antifungal compounds without mycelial contact between the two colonies, and a halo is usually observed), ii) mycoparasitism (the endophyte is able to parasitize pathogen hyphae, impairing their growth), and iii) substrate competition (the endophyte is able to grow more efficiently than the pathogen, without evidence of an inhibition zone).

#### 2.2.2 Volatile antifungal assay

To test the emission of antifungal volatile organic compounds (VOCs) produced by the three fungal endophytes, a VOC assay was performed as described by Chen et al. (2018). Fungal endophytes were cultured independently on petri dishes with YEA medium. Then, the lid was removed and the plate with the endophyte culture was confronted with another plate with an *O. novo-ulmi* culture on MEA medium. In the case of P5 and YM11, a 50 µl drop containing 10 mg ml^−1^ of yeast cells or mycelial fragments was deposited in the middle of the petri dish. YCB36 was subcultured directly using a mycelial plug from the colony edge, which was deposited in the middle of the plate. The two plates (endophyte vs *O. novo-ulmi* and control with no endophyte vs *O. novo-ulmi*) were sealed together with parafilm and incubated vertically at 22 ° C. Three replicates were performed for each endophyte. *O. novo-ulmi* mycelial growth was evaluated at 3, 7 and 10 days by scanning the plates and measuring the colony area using ImageJ free software. Inhibition of *O. novo-ulmi* growth was quantified as described in the previous section.

#### 2.2.3. Antibiotic effect of liquid filtrates against O. novo-ulmi

Endophyte antibiotic potential against *O. novo-ulmi* was evaluated by obtaining liquid filtrates from endophytes. The inoculum source for each liquid culture was obtained by growing each endophyte on YEA medium over an autoclaved cellophane layer. Once grown, endophyte fresh mycelia or yeast cells were harvested, adjusted to 5 mg ml^−1^ with sterile deionized water, and homogenized using an all-glass tissue homogenizer. For each endophyte, 100 µl of the fungal biomass homogenate was added to 70 ml of yeast, malt and glucose liquid medium (4g l^−1^ yeast extract, 10 g l^−1^ malt extract and 4 g l^−1^ glucose; YMG) in Erlenmeyer flasks and grown in the dark at 22 ° C in an orbital shaker (120 rpm). Three replicates were performed for each fungal strain. Glucose content in the liquid media was quantified periodically by the anthrone-sulphuric acid quantification method (Laurentin and Edwards, 2003) using 96-well microplates. The values were quantified according to a standard curve with known concentrations of glucose (0-1000 ppm). Once glucose was totally consumed (at around 25 days of growth), the culture was left to grow for two more weeks to allow secretion of fungal metabolites. After this period, the broth culture was filtered, first with Whatman® filter paper grade 1 (Whatman International Ltd, Maidstone, UK) and subsequently with a sterile 0.2 µm PES syringe filter (Thermo Fisher Scientific, Waltham, MA, USA), to separate broth and fungal biomass. The filtrates were preserved at 4 ° C until use. The antibiotic effect of the liquid filtrates was assessed by growing *O. novo-ulmi* spores in their presence in microplates. To obtain O*. novo-ulmi* spores, mycelial fungal plugs were grown in Erlenmeyer flasks with Tchernoff’s liquid medium (Tchernoff, 1965) at 22 ° C in the dark, under constant shaking to induce sporulation. Three days later spores were collected by centrifugation and adjusted to 10^6^ cells ml^−1^ using a haemocytometer. Six well-replicates were used for each treatment and control group. Microplates were incubated at 22 ° C in the dark and growth was evaluated at 0, 2, 7, 9 and 11 days by reading the optical density (OD) at 630 nm, using a spectrophotometer (ELx808, BioTek, Winooski, VE, USA) (Martín et al., 2010). To evaluate the effect of fungal filtrates, 100 µl of filtrate was mixed with 100 µl of *O. novo-ulmi* spores resuspended in *O. novo-ulmi* liquid filtrate, obtained following the same protocol as for the endophyte liquid filtrates (100 µl of 5 mg ml^−1^ homogenized fresh mycelia grown in YMG for 40 days). Pathogen growth in endophyte liquid filtrates (testing wells) was compared with growth in control wells containing 100 µl of *O. novo-ulmi* spores resuspended in their own liquid filtrate, mixed with a further 100 µl of *O. novo-ulmi* liquid filtrate. Additional background control wells containing endophyte or *O. novo-ulmi* filtrates only (without living cells) were used to subtract the OD of the filtrates from the testing wells, thus measuring only the increased OD due to *O. novo-ulmi* growth. Background control wells also determined that culture filtrates contained no contamination. Comparison between treatment and control groups permitted calculation of *O. novo-ulmi* growth inhibition due to filtrates.

#### 2.2.4. Proteolytic and chitinolytic activities

Extracellular protease and chitinase production by each fungus was evaluated in petri dishes with specific culture media (described below) at 5, 8, 12, 15, 20 and 23 days of incubation. Plugs of the fungal colonies were grown at 22 ° C in the dark. Protease activity was determined on skim milk agar plates (SMA: 28 g l^−1^ skim milk powder, 5 g l^−1^ tryptone, 2.5 g l^−1^ yeast extract, 1 g l^−1^ glucose, 15 g l^−1^ agar; pH 7.0). Positive activity was measured by production of a clear halo surrounding the colonies (Mayerhofer et al., 1973). The halo area (cm^2^) was calculated by subtracting the colony area from the area inside the halo contour surrounding the colony. Chitinase detection medium was prepared as described by Agrawal and Kotasthane (2012). Colloidal chitin was obtained according to Castro et al. (2011). Bromocresol purple dye (Sigma-Aldrich, Darmstadt, Germany) permitted visualization of chitin degradation. A colour shift from yellow to purple around the colony indicates positive chitinase activity. The area showing a colour change was quantified (cm^2^) by subtracting the colony area from the area of the colour-changed halo. Six replicates were performed for each fungus and activity.

#### 2.2.5 Nutrient acquisition mechanisms

Siderophore and HCN production and phosphate solubilization potential were evaluated for each fungus at 5, 8, 12, 15, 20 and 23 days of incubation. Siderophore emission was examined using the Chrome Azurol S (CAS) agar medium. CAS was prepared following the protocol of Schwyn and Neilands (1987), in which 60.5 mg CAS (Merck KGaA, Darmstadt, Germany) was dissolved in dH_2_O and mixed with 10 ml of iron (III) solution (1mM FeCl_3_.6H_2_0, 10 mM HCl). After this, hexadecyltrimethylammonium bromide (HDTMA; Sigma-Aldrich, Darmstadt, Germany) was dissolved in water, mixed with CAS solution and autoclaved at 121 ° C. The basal medium was prepared using 0.5 % succinic acid, 0.4 % KH_2_PO_4_, (NH_4_)_2_SO_4_ and 2 % agar at pH 5.3. The HDTMA-CAS solution was slowly mixed until the desired blue colour was obtained. Plugs of fungal colonies were grown in six replicate petri dishes containing HDTMA-CAS agar medium at 22 ° C in the dark. Positive siderophore activity was evaluated by colour change from blue to yellow, subtracting the colony area from the area of the colour-changed halo (cm^2^).

The hydrogen cyanide (HCN) production test was performed based on Millar and Higgins (1970). HCN solution was prepared by mixing 0.5% picric acid and 2% sodium carbonate. Sterilized Whatman® filter papers were soaked in HCN solution and placed on the lid of YEA petri dishes where fungal plugs were cultured. Then, petri dishes were sealed with parafilm and incubated in the dark at 22 ° C. Six replicates were performed for each fungus. Colour change in the filter paper indicates positive HCN release by the fungi, due to the ability of HCN to reduce picric acid to isopurpuric acid. Activity was scored as negative (no colour change) or positive (brown to reddish brown).

Phosphate solubilization was performed in Pikovskaya agar medium with the addition of 0.3 % insoluble calcium triphosphate (Sigma-Algrich, Darmstadt, Germany). Endophyte plugs were grown in six replicate petri dishes with Pikovskaya agar medium at 22 ° C in the dark. The presence of a halo or clear area around the colony, indicating phosphate solubilization potential, was measured as halo area (cm^2^).

#### 2.2.6. Nutrient utilization patterns and chemical sensitivity

Nutrient utilization patterns and the sensitivity of the endophytes and the pathogen to a series of inhibitory substrates were evaluated using in-house configured phenotypic microarrays, following the methodology described by Blumenstein et al. (2015). Fungal inocula were prepared as described in the section “Antibiotic effect of liquid filtrates against *O. novo-ulmi*”, but in this case the fungal biomass was resuspended in Murashige and Skoog (MS) liquid medium. Fungal growth on different nutrients was evaluated in 96-well microplates by measuring OD at 630 nm using an ELx808 spectrophotometer after 9 days of incubation. The following carbon sources were added individually to the MS medium: glucose, fructose, sucrose, cellobiose, pectin, xylose, maltose, starch and carboxymethyl-cellulose. All nutrient solutions were adjusted to a final concentration of 0.25 M carbon atoms. Various metabolites with potential to inhibit fungal development were added individually to the MS medium: gallic acid (1 g l^−1^), salicylic acid (0.05 g l^−1^), tannic acid (1 g l^−1^), veratryl alcohol (2.37 g l^−1^), quercetin (0.01 g l^−1^), catechin (1 g l^−1^), thymol (0.05 g l^−1^) and ethanol (5%). Carbon consumption by endophytes was compared with carbon consumption by *O. novo-ulmi* using the Niche Overlap Index (NOI). NOI was calculated as the number of carbon sources consumed by the endophyte and the pathogen, divided by the number of sources consumed by the pathogen (Lee and Magan, 1999). A NOI value of 0.9 or above indicates a high degree of niche overlap and a competitive disadvantage for the pathogen (Blumenstein et al., 2015).

#### 2.2.7. Phytohormone production by endophytes and the pathogen

Ability of the three endophytes and *O. novo-ulmi* to produce auxins and gibberellins was evaluated. For indole-3-acetic acid (IAA) detection, the four fungi were grown separately in 2% malt extract broth (MEB) containing 0.2% (v/v) L-tryptophan. Erlenmeyer flasks contained 30 ml medium and 60 µl homogenized fungal biomass adjusted to 5 mg ml^−1^. Flasks were kept in the dark at 22 ° C under constant shaking for 15 days. IAA production was estimated following the colorimetric protocol, using Salkowski reagent described in Gang et al. (2019) prepared by mixing 0.5 M ferric chloride (FeCl_3_) and 35% perchloric acid (HClO_4_). Three replicates were performed for each fungus. To quantify the IAA produced by each fungus, three aliquots of each broth were selected and centrifuged at 10,000 rpm for 10 min. After that, 500 µl supernatant was mixed with 1 ml Salkowski reagent and incubated for 25 min in the dark. Absorbance was read at 530 nm, and the concentration of IAA produced by each isolate was determined using a standard curve of pure IAA (Sigma-Aldrich, Darmstadt, Germany) ranging from 0 to 1000 µg ml^−1^.

Gibberellic acid (GA_3_) released by each fungus was quantified using the liquid filtrates obtained for the antibiotic evaluation (see section 2.2.3). GA_3_ quantification was performed by selecting three aliquots of the filtrate and adjusting the pH to 1.5 using 0.1 M HCl. Then, 2 ml of the acidified liquid medium was mixed with 4 ml ethyl acetate, vortexed vigorously and centrifuged at 5,000 rpm for 10 min (Holbrook et al., 1961). This process was repeated two more times, after which the organic phase was mixed, evaporated and suspended in absolute ethanol. Absorbance was read at 254 nm and recorded at 20 s intervals for 2 min according to Berríos et al. (2004). GA_3_ concentration was estimated using GA_3_ (Sigma-Aldrich, Darmstadt, Germany) standard solutions in ethanol ranging from 0.1 g l^−1^ to 1 g l^−1^.

### 2.3 In vivo experiment

#### 2.3.1 Plant material

The plant material used in the *in vivo* experiment was produced in 2012 as part of the regular activities of the Spanish elm breeding programme at “Puerta de Hierro” Forest Breeding Centre. Controlled crosses were performed between two *U. minor* genotypes (M-CC1 and AL-AN1). To properly evaluate offspring susceptibility to DED, the seedlings obtained (N=37) were cloned through aerial cuttings. After the susceptibility tests, four seedlings showing moderate to high DED susceptibility (50-70 % leaf wilting 60 days after *O. novo-ulmi* inoculation) were selected and 30 to 34 clonal replicates of each seedling were planted in an experimental plot in 2014 (Table 1). The plot had a complete random design and plants were spaced 1 × 1.4 m apart. Plants were regularly watered in spring and summer to avoid water stress. One highly DED resistant commercial clone (Sapporo, N=8) and one highly DED susceptible clone (M-PZ3, N=8) were also included in the plot to confirm the virulence of the *O. novo-ulmi* isolate.

**Table 1:** Fungal treatment and plant material specifications. E-O-: control plants without endophyte and *O. novo-ulmi*; E+O-: endophyte-treated plants without *O. novo-ulmi*; E-O+: *O. novo-ulmi*-treated plants without previous endophyte treatment; E+O+: endophyte- and *O. novo-ulmi-*treated plants.

| Inoculated endophytes | Plant genotype | DED-tolerance level<sup>1</sup> | Treatment | N |
| --- | --- | --- | --- | --- |
| P5+YM11 | (M-CC1 × AL-AN1).5 | Susceptible | E-O- | 5 |
|  |  |  | E+O- | 5 |
|  |  |  | E-O+ | 10 |
|  |  |  | E+O+ | 10 |
|  | (M-CC1 × AL-AN1).7 | Susceptible | E-O- | 5 |
|  |  |  | E+O- | 5 |
|  |  |  | E-O+ | 10 |
|  |  |  | E+O+ | 10 |
|  | (M-CC1 × AL-AN1).8 | Susceptible | E-O- | 5 |
|  |  |  | E+O- | 5 |
|  |  |  | E-O+ | 10 |
|  |  |  | E+O+ | 10 |
| YCB36 | (M-CC1 × AL-AN1).15 | Susceptible | E-O- | 6 |
|  |  |  | E+O- | 12 |
|  |  |  | E-O+ | 11 |
|  |  |  | E+O+ | 5 |
| --- | Sapporo | Highly tolerant | E-O- | 8 |
|  |  |  | E-O+ | 8 |
| --- | M-PZ3 | Highly susceptible | E-O- | 8 |
|  |  |  | E-O+ | 8 |

<sup>1</sup> According to previous susceptibility tests

#### 2.3.2 Experimental design

*In planta* inoculations were performed in May 2017, when plants were 6-years old. Two endophyte cell suspensions were prepared (as described in the next section): (i) a mixture of P5 and YM11; and ii) YCB36. These suspensions were inoculated into the vascular system in the middle of the trunk using an endotherapy device (BITE, Padova, Italy). *Ophiostoma novo-ulmi* inoculations were performed with a bud-cell suspension (prepared as described in the next section) 5 cm below the endophyte inoculation point, on the opposite side of the trunk. Pathogen spores were delivered into the xylem sap stream through a transverse cut made with a sharp blade (Solla et al., 2005). Four treatments were applied (Fig. S1): (i) Control plants with no endophyte or pathogen inoculation (E-O-); (ii) Plants inoculated with endophytes (P5+YM11 or YCB36) without subsequent *O. novo-ulmi* inoculation (E+O-); iii) Plants inoculated with *O. novo-ulmi* without previous endophyte inoculation (E-O+); and iv) Plants inoculated with endophytes subsequently inoculated with the pathogen (E+O+) (Table 1). In treatments i) to iii), plants not inoculated with either the endophytes or the pathogen (E- and O- treatments) received sterile distilled water. Endophyte and *O. novo-ulmi* inoculations, and their corresponding control treatments, were performed on 1st May and 16th May, respectively (Fig. S1).

#### 2.3.3. Inoculum preparation

To obtain the endophyte inoculum, a liquid suspension of bud cells (for YM11 and P5) or hyphal fragments (for YCB36) were prepared. Endophytes were grown in petri dishes containing YEA medium, with (YM11 and YCB36) or without (P5) an autoclaved cellophane layer, for 10 days at 22 ° C in the dark. P5 yeast cells were scraped from the agar using a sterile spatula and suspended in sterile distilled water. YM11 formed a colony of dense, hard agglomerates of bud cells, requiring the use of an all-glass tissue homogenizer to obtain a homogeneous cell suspension. YCB36 did not produce a sufficient quantity of conidia and therefore spores were replaced by mycelial fragments also obtained using an all-glass tissue homogenizer, first with a large clearance pestle and then with a small one (∼50 strokes with each). We then poured the resulting suspension over sterilized cotton wool, so the longer fungal hyphae were collected into the cotton wool. The resulting suspension was quite homogenous, with hyphal fragments of 1-4 cells/fragment. Aqueous suspensions of yeast cells or mycelial fragments were adjusted using a haematocytometer to obtain a final inoculum concentration of 2 x 10^6^ cells ml^−1^ for the mixture of P5 and YM11 (10^6^ cells ml^−1^ of each fungus) and approximately 10^6^ cells ml^−1^ for YCB36.

To obtain *O. novo-ulmi* spores, mycelial fungal plugs were grown in Erlenmeyer flasks with Tchernoff’s liquid medium (Tchernoff, 1965) at 22 ° C in the dark under constant shaking to induce sporulation. Three days later, spores were collected by centrifugation and adjusted to 10^6^ blastospores ml^−1^ using a haemocytometer.

#### 2.3.4. Leaf wilting evaluation

Leaf yellowing and wilting are typical DED symptoms. As the disease progresses, a mixture of healthy and suffering foliage may be seen (Brasier and Buck, 2001) and the percentage of foliar wilting in the crown is a good indicator of tree decline (Solla et al., 2005). Leaf wilting for symptoms of *O. novo-ulmi* was evaluated in each tree at 30, 60 and 120 days post inoculation (dpi) and also 1 year post-inoculation (ypi). Assessments were performed by three independent evaluators and the mean of the three scores was calculated for each tree and treatment. A representative leaf of the average wilting status of the tree was selected to measure leaf stomatal conductance. This measurement was made at the time of *O. novo-ulmi* inoculation and at 30 dpi using an SC-1 leaf porometer (Decagon Devices, Washington, USA) in five trees per treatment, at midday on both occasions.

#### 2.3.5. In planta endophyte detection

To evaluate plant colonization by endophytes, wood samples from control (E-O-) and endophyte-inoculated trees (E+O-) were extracted using an increment borer at 75 dpi and 1 ypi. Two samples per tree were taken, at 5 and 50 cm above the endophyte inoculation point. Three replicate trees per treatment were evaluated. Fungal DNA presence was quantified through quantitative PCR (qPCR). First, core wood samples were ground to a fine powder in a ball mill (Mixer mill MM 400, Retsch GmbH, Haan, Germany) and DNA was extracted using the Norgen plant/fungi DNA isolation kit (Norgen Biotek Corp., Thorold, ON, Canada), obtaining high yield and quality scores. Primer sequences were designed within the sequences obtained by ITS Sanger sequencing of each fungus (described previously) using Primer3 Version 0.4.0 (http://bioinfo.ut.ee/primer3-0.4.0/primer3/) and following specific parameters for qPCR primer design (Udvardi et al., 2008). Specificity tests for the primers designed were performed by selecting 10 additional fungi available in our collection and amplifying fungal DNA through conventional PCR. If an amplification product was observed, the primer pair was discarded and new primer pairs were designed. Selected primer pairs were set as follows: the primer pair for P5 (forward primer: 5’-CAACGGATCTCTTGGCTCTC-3’/ reverse primer: 5’- AACAGACATACTCTTCGGAATACC-3’) created a 136 base-pair (bp) product, the primer pair for YM11 (forward primer: 5’- AGGAACTGGCCTCAAAGACA-3’/ reverse primer: 5’- TCCTACCTGATCCGAGGTCA -3’) created a 142 bp product, and the primer pair for YCB36 (forward primer: 5’-AGTATACGCCGCCTTGACAC-3’/ reverse primer: 5’- GTCGTAAAACATGGGGAACG-3’) created a 130 bp product. Relative fungal DNA estimations were performed independently for each fungus using SSoFast EvaGreen® Supermix (Bio-Rad Laboratories, Irivine, CA, USA) in a CFX96 real-time PCR detection system thermocycler (Bio-Rad Laboratories, Irvine, CA, USA). Fungal DNA was standardized per quantity of plant DNA by amplifying a fragment of the *U. minor* ITS region (ITSUlmi) (forward primer: 5’- ATATGTCAAAACGACTCTCGGCAAC-3’/ reverse primer: 5’-AACTTGCGTTCAAAGACTCGATGGT-3’). In triplicate, reactions contained 2 μl sample DNA and 8 μl master mix. Following a first denaturation step at 95 ° C for 30 s, 40 cycles were performed at 95 °C for 5 s and 60 °C for 30 s. Each plate included two replicates of each sample: one for amplification of the specific fungal ITS, and the other for amplification of *U. minor* ITS. Three no-template controls were included for each primer pair. The amplification results were analysed and expressed as fungal presence relative to non-inoculated plants.

#### 2.3.6. Plant biochemical measurements

Wood samples were extracted from trees showing a significant effect of the endophyte inoculation treatment on plant symptomatology, to measure non-structural carbohydrates, total phenolic and flavonoid content, and proline. The concentration of non-structural carbohydrates (NSC) was measured using a modified protocol from Maness (2010). Soluble sugars (SS) were extracted by incubating 50 mg fresh material in 1 ml of 80% ethanol. An aliquot of the ethanol extract was reserved for proline quantification (see below). The insoluble material in the pellet was used for starch determination. Sugar monomers from the ethanol extract or starch digestion were quantified by the anthrone-sulphuric acid colorimetric microassay, based on Laurentin and Edwards (2003), using 96-well microplates. The values were quantified according to standard curves with known concentrations of glucose, fructose and galactose (0- 1000 ppm) in the case of SS, and only glucose (0-1000 ppm) in the case of starch. Total NSC concentration was calculated by adding the concentrations of SS and starch.

Total phenolic content extraction was performed according to a microplate-adapted protocol described by Ainsworth and Gillespie (2007) and quantified using the Folin-Ciocalteu reagent (F-C). Briefly, 20 mg powdered wood was extracted in 1 ml of 95 % methanol under constant shaking in a mixer mill (Precellys, Bertin Instruments, Montigny-le-Bretonneux, France), then incubated at room temperature for 48 h in the dark. The supernatant was recovered and 200 µl of the F-C reagent at 10 % (v/v) was added per each 100 µl of extract or standard. Finally, 800 µl of Na_2_CO_3_ 700 mM was added and incubated at room temperature for 2 h. Absorbance was read at 765 nm and the values were quantified according to a standard curve with known concentrations of gallic acid (25 µM-1.5 mM). Results were expressed as mg equivalent gallic acid per gram of fresh weight of sample.

Total flavonoid content was determined using the same extract obtained for total phenolic content and quantified by the colorimetric method described in Barreira et al. (2008), with slight modifications. Briefly, 177.5 μL aliquots of each sample and 25 μL aliquots of each standard point (diluted with 152.5 μL deionized water) were mixed with NaNO_2_ (5%, 7.5 μL). After 6 min, 15 μL of a 10% AlCl_3_·6H_2_O solution was added, and after 5 min incubation at room temperature, 50 μL of 1M NaOH was added. The absorbance of each blank, consisting of the same sample mixtures but with deionized water instead of 10 % AlCl_3_·6H_2_O solution, was subtracted from the test absorbance. The absorbance was read at 510 nm using a microplate reader and the values were quantified according to a standard curve with known concentrations of quercetin (0.2-1.0 mg ml^−1^ in ethanol). Results were expressed as mg equivalent quercetin per gram of fresh weight of sample.

Free proline content was determined with the same extract used for SS determination. Quantification was carried out using a microplate reader protocol (Carillo and Gibon, 2011). In each well, 50 µl of the ethanolic extract, blank or proline standards (1-0.4-0.2-0.1-0.04 mM) was mixed with 100 µl of the reaction mix: 1 % ninhydrin (w/v) in 60 % acetic acid (v/v) and 20 % ethanol (v/v). Plates were incubated in a water bath at 95 ° C for 20 min and absorbance was read at 520 nm. The readings were transformed to proline contents (nmol mg^−1^ FW) according to Carillo and Gibon (2011).

### 2.4 Statistical analysis

The effect of endophytes on *O. novo-ulmi* growth in *in vitro* assays was tested with one-way ANOVA and the post-hoc Fisher’s Least Significant Difference (LSD) test (*P*<0.05). The area of the halo formed by the four fungal strains was compared using a one-way ANOVA test followed by Fisher’s LSD test (*P*<0.05) to identify differences in protease, chitinase and siderophore emission between isolates. The effect of inhibitory substances and carbon sources on fungal growth was represented with a two-way joining cluster analysis using the heatmap function in the “pheatmap” package in R-studio (R Core Team 2017). Euclidean distances were used in the clustering of fungal endophytes or chemicals. Differences in wilting symptoms, leaf stomatal conductance and biochemical variables among treatments were analysed with one-way ANOVA test followed by Fisher’s LSD test (*P*<0.05). To observe differences in wilting symptoms among elm clones due to *O. novo-ulmi*, a factorial ANOVA was performed, taking the clone and *O. novo-ulmi* treatments as factors. Fungal quantification through quantitative PCR was performed for each sample at 5 and 50 cm from the inoculation point with three technical replicates per PCR plate. The cycle threshold value (Ct) obtained for the plant and endophyte ITS in each sample was transformed to double delta Ct (ΔΔCt). Then, ΔΔCt of the endophyte was normalized to ΔΔCt of plant ITS. The normalized values of endophyte-inoculated plants were relativized to control plants, obtaining the fold-change of endophyte presence. These values were analysed with a one-way ANOVA test to determine whether there were significant differences between inoculated and non-inoculated trees.

Analyses were run using STATISTICA version 8.0 (StatSoft, Tulsa, OK, USA) and R software (R Core Team 2017). In all cases, normality was confirmed before ANOVA analysis with skewness and kurtosis values and the Kolmogorov-Smirnov test.

## 3. Results

### 3.1. Molecular identification of fungal endophytes

According to the top BLAST hits for ITS and LSU sequencing the P5 endophyte is assigned to the genus *Cystobasidium* (Cystobasidiomycetes) and YM11 to *Exophiala* (Eurotiomycetes) (Table 2). However, the top BLAST hit for *TEF1α* sequence of YM11 assigned it to *Knufia*, although with lower percentage of identity than ITS and LSU sequences (Table 2). Results for YCB36 indicate that this fungus belongs to the Phaeosphaeriaceae family (Dothideomycetes). According to the LSU sequence the top hit corresponded to the genus *Hydeomyces* (99.41% identity), while the rest of the markers did not provide significant coverages and/or identities at the genus level (Table 2). Therefore, it might be an undescribed fungus within Phaeosphaeriaceae.

**Table 2:**
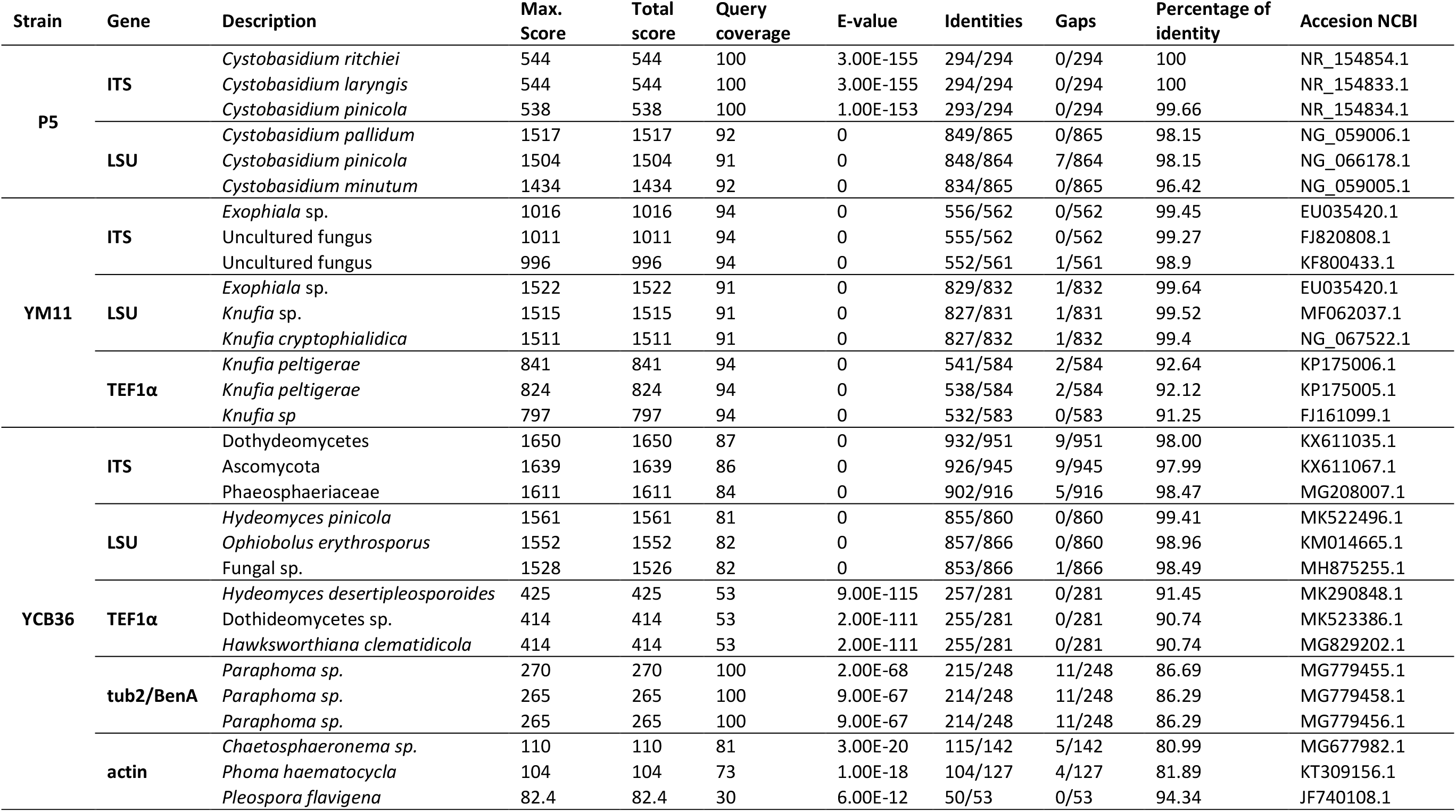
Top three BLAST hits based on nucleotide megablast of ITS and LSU of rDNA sequences and *TEF1α*, *β-tub* or actin as protein coding genes. The corresponding GenBank taxa identity and NCBI accession numbers are indicated.

### 3.2. In vitro fungal endophyte characterization

#### 3.2.1. In vitro endophyte-pathogen interactions

*O. novo-ulmi* growth was reduced when confronted with YCB36 and P5 endophytes, but not with YM11 despite this endophyte displaying some visible slight inhibition of *O. novo-ulmi* (Table 3; Fig. S2A, D). When confronted with YCB36, *O. novo-ulmi* growth was reduced between 28.05 ± 3.59 % and 37.62 ± 5.32 % throughout the experiment (Table 3; Fig. S2A, D). Volatiles emitted by YCB36 and P5 also inhibited *O. novo-ulmi* growth (Table 3). The highest inhibition was observed with YCB36 (32.82 ± 5.54 %) and a moderate inhibition was observed with P5 (11.02 ± 3.28 %) (Table 3; Fig. S2B, D).

**Table 3:** Influence of fungal endophytes on *O. novo-ulmi* growth in the dual culture assay (day 12 of incubation), volatiles (VOCs) assay (day 10) and antibiosis assay (day 10). Different letters in the same row indicate significant differences between fungi, according to Fisher’s post-hoc test (*P*<0.05).

| Assay | Units | *O. novo-ulmi* growth |  |  |  |
| --- | --- | --- | --- | --- | --- |
|  |  | Control | YCB36 | P5 | YM11 |
| Dual culture | cm<sup>2</sup> | 25.567 ± 3.69<sup>c</sup> | 15.48 ± 1.03<sup>a</sup> | 18.95 ± 0.48<sup>b</sup> | 25.54 ± 1.72<sup>c</sup> |
| VOCs | cm<sup>2</sup> | 30.85 ± 3.27<sup>c</sup> | 19.98 ± 1.64<sup>a</sup> | 24.67 ± 1.92<sup>b</sup> | 29.29 ± 1.46<sup>c</sup> |
| Antibiosis | OD | 0.150 ± 0.001<sup>c</sup> | 0.062 ± 0.003<sup>a</sup> | 0.134 ± 0.011<sup>b</sup> | 0.101 ± 0.008<sup>b</sup> |

#### 3.2.2. Antibiotic traits

*O. novo-ulmi* growth was reduced when cultured in liquid media containing YM11 and YCB36 liquid filtrates (Table 3; Fig. S2C). YCB36 filtrate was able to reduce *O. novo-ulmi* growth by 55.94 ± 0.96 % (*P*<0.01; Table 3; Fig. S2C), while YM11 filtrate reduced *O. novo-ulmi* growth by 28.74 ± 2.9 % (Fig. S2C). P5 filtrate did not reduce *O. novo-ulmi* growth (Table 3; Fig. S2C). Proteolytic activity varied considerably among isolates, with *O. novo-ulmi* showing a halo formation of 28.73 cm^2^ (Table 4, Fig. S3). Endophyte P5 had low proteolytic activity, and the halo area was barely visible (1.33 ± 0.06 cm^2^). In contrast, YCB36 and YM11 showed a clear halo around the colony, which was significantly greater for YCB36 than for YM11 (10.28 ± 2.48 and 5.75 ± 0.71 cm^2^, respectively) (Table 4; Fig. S3). In the case of chitinolytic activity, YCB36 produced a clear blue-purple halo surrounding the colony (18.19 cm^2^ at day 5 and 49.85 ± 1.63 cm^2^ at day 20), while the other endophytic fungi had no chitin degradation ability (Table 4, Fig. S3). Chitinase activity in *O. novo-ulmi* became visible at day 20 (6.46 ± 1.96 cm^2^) (Table 4, Fig. S3).

**Table 4:** Protease and chitinase activities, siderophore and hydrogen cyanide production, and phosphate solubilisation in *in vitro* assays of the fungal endophytes and the pathogen at the end of the experiment (day 20). Different letters in the same row indicate significant differences between fungi, according to Fisher’s post-hoc test (*P*<0.05).

| Activity | Units | Fungal strain |  |  |  |
| --- | --- | --- | --- | --- | --- |
|  |  | *O. novo-ulmi* | YCB36 | P5 | YM11 |
| Protease | cm<sup>2</sup> (halo) | 28.73 ± 1.96<sup>c</sup> | 10.28 ± 2.48<sup>b</sup> | 1.33 ± 0.06<sup>a</sup> | 5.75 ± 0.71<sup>b</sup> |
| Chitinase | cm<sup>2</sup> (halo) | 6.46 ± 1.93<sup>a</sup> | 49.85 ± 1.63<sup>b</sup> | negative | negative |
| Siderophore | cm<sup>2</sup> (halo) | 5.39 ± 0.98<sup>a</sup> | 7.90 ± 0.44<sup>b</sup> | negative | negative |
| Hydrogen cyanide | positive/negative | negative | negative | negative | negative |
| Phosphate solubilization | cm<sup>2</sup> (halo) | negative | negative | negative | negative |

#### 3.2.3. Nutrient acquisition mechanisms and nutritional niche

Only the *O. novo-ulmi* and YCB36 isolates secreted siderophores into the medium (Table 4, Fig. S3). YCB36 exhibited greater halo formation than *O. novo-ulmi* at day 20 (7.90 ± 0.44 vs 5.39 ± 0.98 cm^2^) (*P*<0.05) (Table 4, Fig. S3). However, neither HCN nor phosphate were solubilized by any of the fungal strains evaluated (Table 4).

The carbon utilization pattern and chemical sensitivity to inhibitory substances of each isolate are shown in the two-way joining cluster graph (Fig. 1). As expected, the analysis of the substrates (vertical axis) provided two clusters: the carbon sources on one side and the inhibitory metabolites on the other. The clustering of the fungal isolates (horizontal axis) separated P5 from the other fungi, coinciding with the taxonomic division of the fungi into Basidiomycota and Ascomycota, respectively. The clustering also separated the two ascomycetous endophytes (YM11 and YCB36) from the pathogen. P5 and *O. novo-ulmi* were less affected by inhibitory metabolites (e.g., by gallic acid and thymol), while YM11 and *O. novo-ulmi* showed the highest growth in the presence of certain carbon sources (e.g., sucrose, glucose and fructose).

**Fig. 1:**
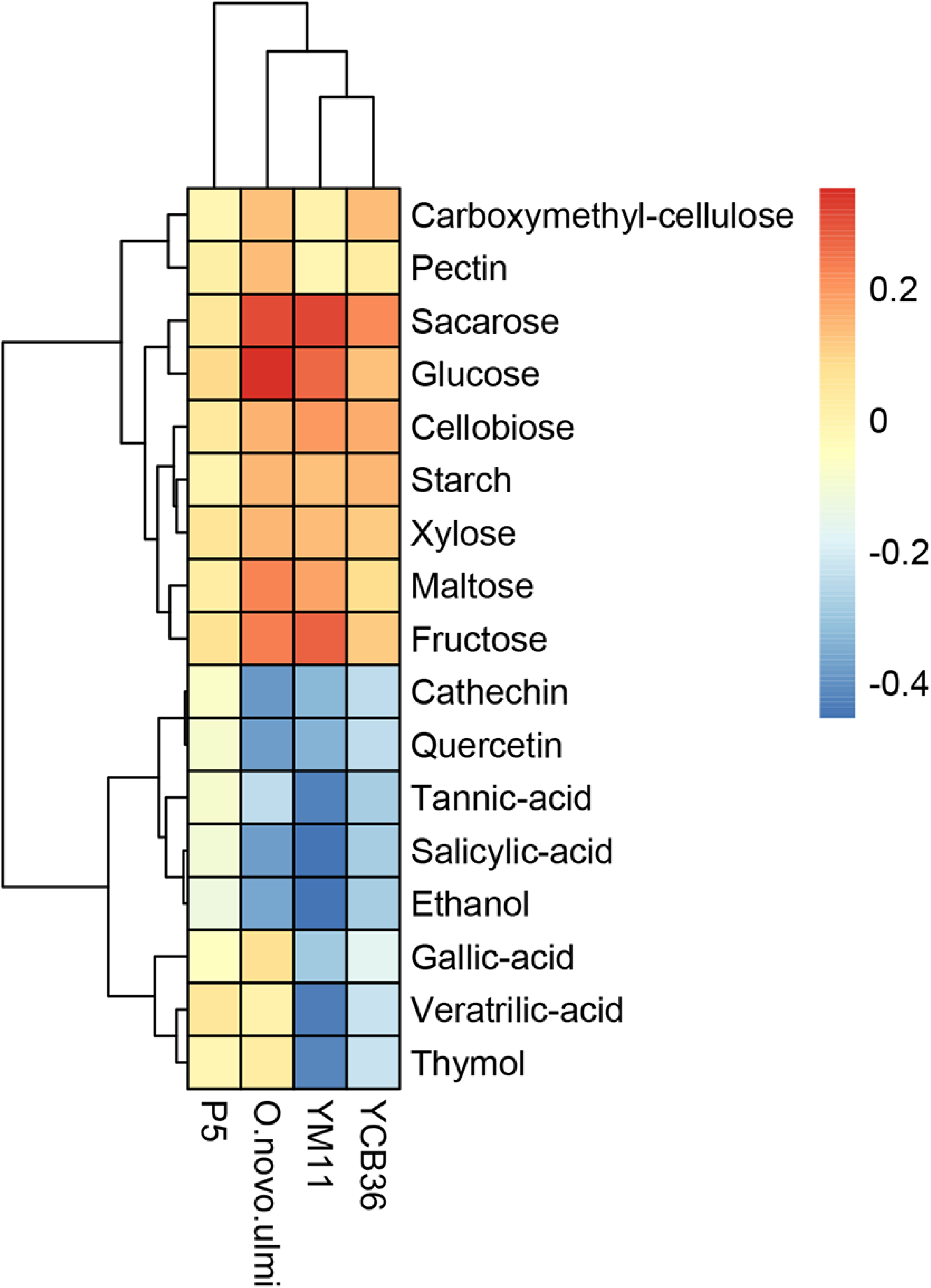
Two-joining clustering analysis of the in-house configured phenotypic arrays of fungal endophytes and *O. novo-ulmi* (horizontal axis) growing with different nutrients and inhibitory substrates (vertical axis). Euclidean distances were used in the clustering of fungal endophytes and substrates. The colour legend indicates inhibition (bluish colours) or promotion (reddish colours) of fungal growth.

Only considering carbon sources, the NOI of *O. novo-ulmi* with regard to P5, YM11 and YCB36 was 0.78, 0.78 and 1, respectively, indicating a putative high niche overlap between the pathogen and YCB36.

#### 3.2.4. Growth promotion potential

The three endophytes and the pathogen produced IAA. *O. novo-ulmi* produced 191.23 µg IAA g^−1^ fungal dry weight. P5 produced 1110.14 µg IAA g^−1^ fungal dry weight and was the endophyte with the highest IAA production (Fig. 2A). GA_3_ production was similar in *O. novo-ulmi*, YCB36 and P5 (0.196 ± 0.014 g l^−1^, 0.258 ± 0.010 g l^−1^ and 0.173 ± 0.020 g l^−1^, respectively), while YM11 produced the lowest GA_3_ concentration (0.108 ± 0.042 g l^−1^) (Fig. 2B).

**Fig. 2:**
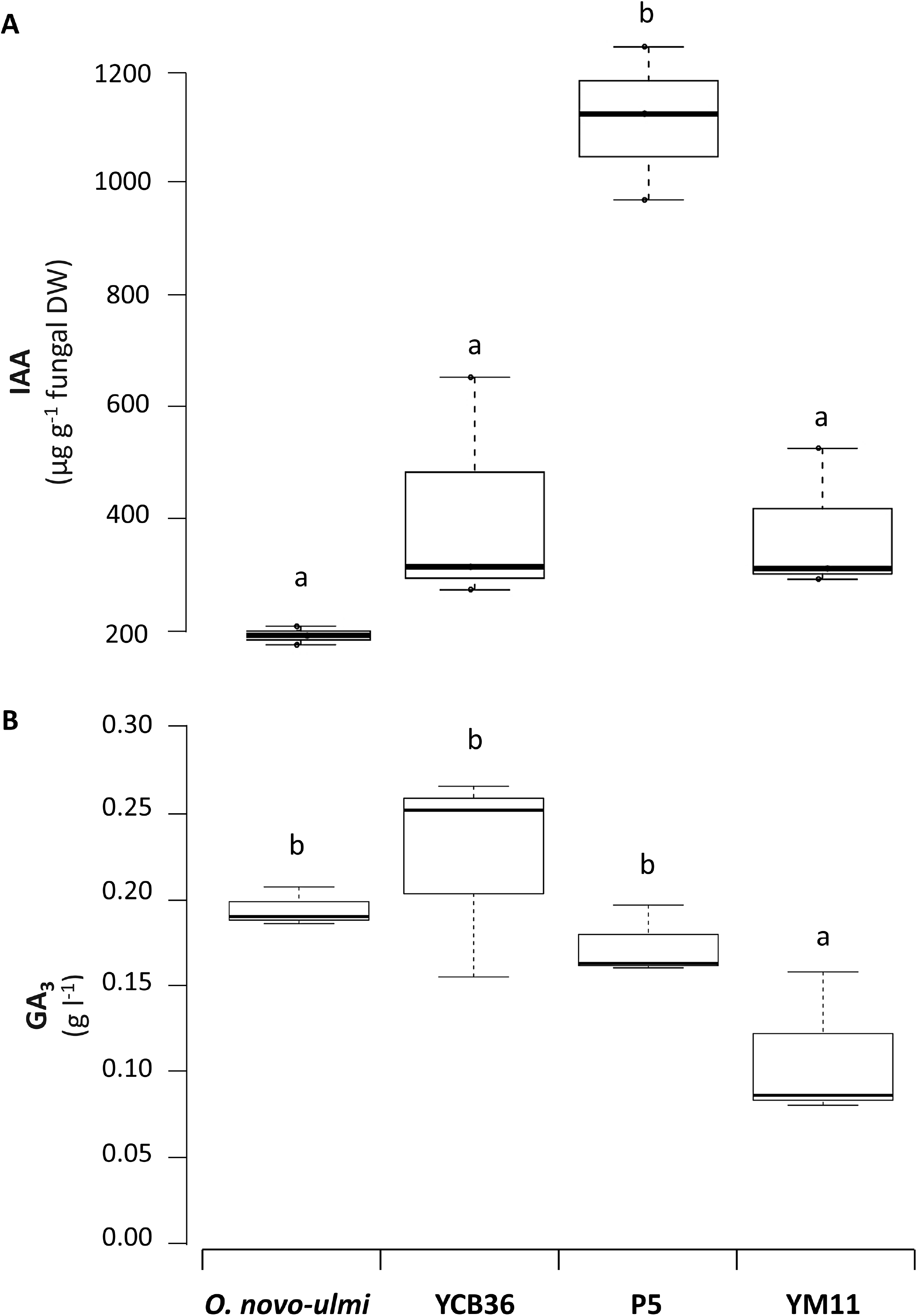
Indole-3-acetic acid (IAA) and gibberellic acid (GA_3_) production by fungal endophytes and pathogen**. (A)** IAA production by *O. novo-ulmi* and fungal endophytes (YCB36, P5 and YM11) after 15 days of incubation in malt extract broth containing 0.2 % L-tryptophan. **(B)** GA_3_ content in liquid filtrates after which *O. novo-ulmi* and the three endophytes have been growing in YMG for 40 days. Different letters indicate significant differences between fungi according to Fisher’s post-hoc test (P<0.05).

### 3.3. In vivo assay

#### 3.3.1. Leaf wilting evaluation and stomatal conductance

Leaf wilting at 120 dpi in the resistant control clone Sapporo was low (5.94 % ± 2.59; mean ± SE), but high in the susceptible control clone M-PZ3 (68.33 % ± 11.76), confirming the virulence of the *O. novo-ulmi* isolate inoculated. Trees that were not inoculated with *O. novo-ulmi* (E-O- and E+O-) showed no leaf yellowing or wilting symptoms, while trees inoculated with the pathogen (E-O+ and E+O+) displayed various levels of leaf wilting. The four genotypes of the progeny (M-CC1 × AL-AN1) exhibited a similar response to *O. novo-ulmi* (*P* = 0.423). Therefore, the effect of the genotype was not considered in subsequent analyses. Trees not inoculated with endophytes (E-O+) showed mean leaf wilting values of 48.93 to 66.95% at different evaluation dates (Fig. 3A,C). Trees inoculated with P5+YM11 before *O. novo-ulmi* inoculation (E+O+) showed similar symptoms to non-inoculated trees (E-O+) (Fig. 3A). In this case, the proportion of trees with leaf wilting percentages higher than 50% was similar in E-O+ and E+O+ plants at 120 dpi (Fig. 3B). However, trees inoculated with YCB36 (E+O+) showed lower wilting symptoms than non-inoculated trees (E-O+) at 30, 60 and 120 dpi (Fig. 3C). In this case, the proportion of trees with leaf wilting percentages higher than 50% was lower in E+O+ trees than in E-O+ trees at 120 dpi (Fig. 3D). After 1 year, however, the protective effect of the YCB36 isolate was no longer evident (Fig. 3C).

**Fig. 3:**
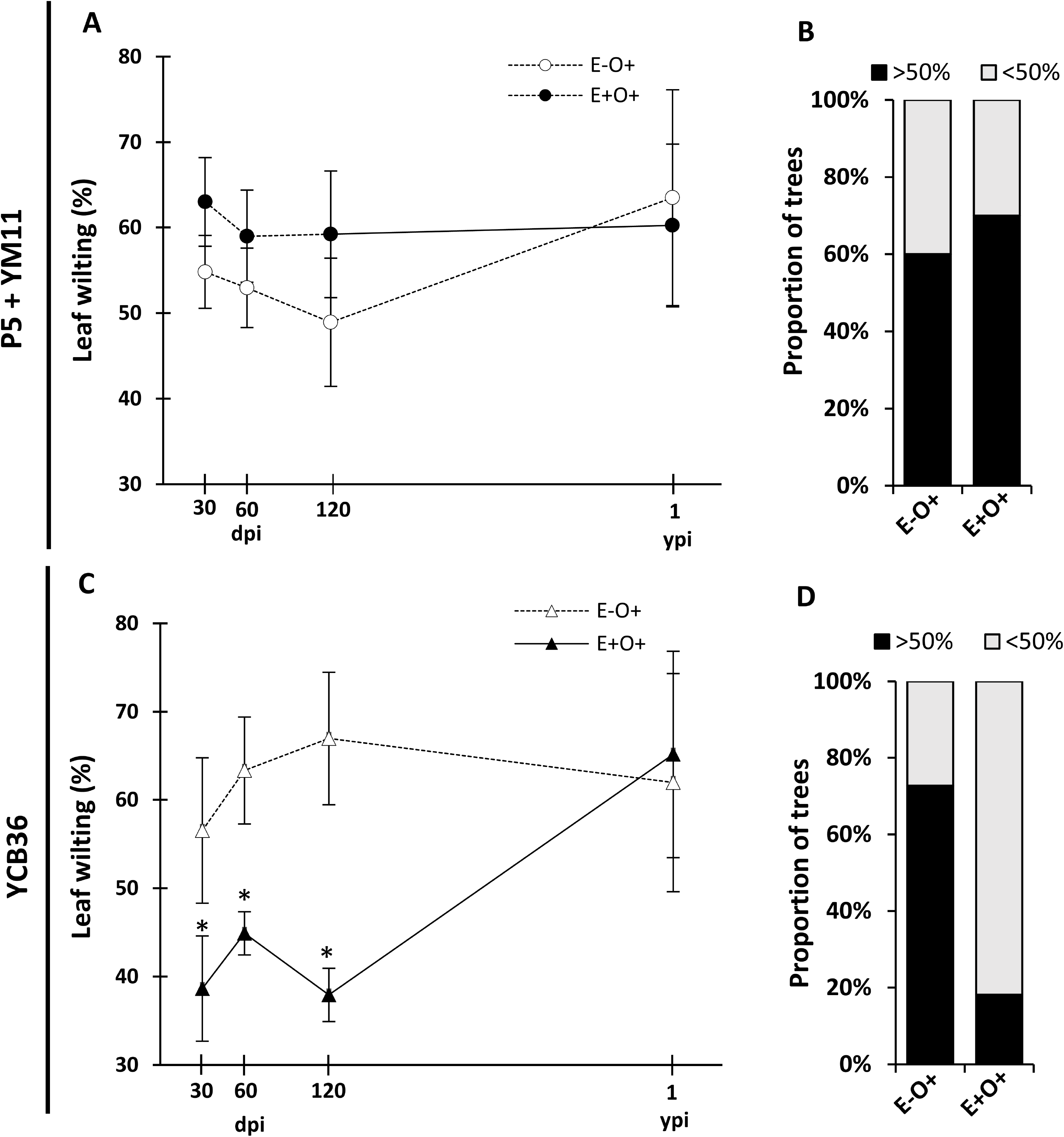
Leaf wilting induced by *O. novo-ulmi* inoculation at 30, 60, 120 dpi and 1 ypi in: P5+YM11-inoculated trees **(A),** and YCB36-inoculated trees **(C).** The proportion of trees with leaf wilting percentages higher or lower than 50% is shown in (**B)** for P5+YM11-inoculation and in (**D)** for YCB36-inoculation. E-O+ indicates non-endophyte-inoculated trees with *O. novo-ulmi* inoculation, and E+O+ indicates endophyte-inoculated trees with *O. novo-ulmi* inoculation. Non- *O. novo-ulmi*-inoculated trees (not shown) showed no wilting symptoms. Asterisks in graphs (A) and (B) indicate significant differences between treatments according to Fisher’s post-hoc test (P<0.05).

In addition, *O. novo-ulmi* inoculation caused a decline in stomatal conductance at 30 dpi, both in trees pre-inoculated with P5+YM11 (E+O+) (Fig. 4A, B) and E-O+ trees (Fig. 4a,b). YCB36-inoculated trees (E+O-, Fig 4B) showed higher stomatal conductance at 30 dpi than non-inoculated trees (E-O-, Fig. 4B).

**Fig. 4:**
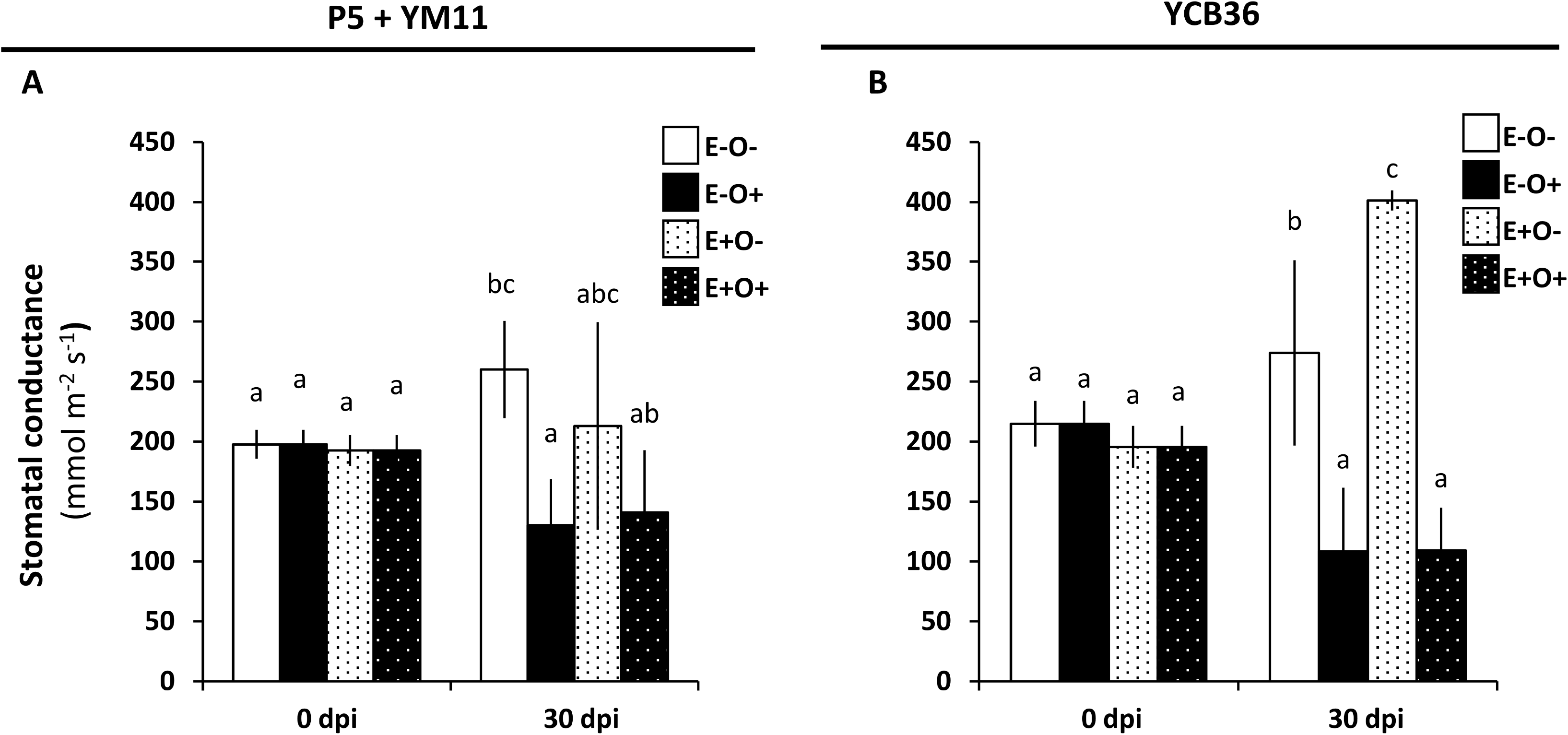
Stomatal conductance measured at 0 dpi and 30 dpi in P5 + YM11-inoculated plants **(A)** and YCB36-inoculated plants **(B).** Treatments were: trees without endophyte or *O. novo-ulmi* inoculation (E-O-); non-endophyte-inoculated trees with *O. novo-ulmi* inoculation (E-O+); endophyte-inoculated trees without *O. novo-ulmi* inoculation (E+O-), and trees inoculated with endophyte before *O. novo-ulmi* inoculation (E+O+). Different letters indicate significant differences between treatments according to Fisher’s post-hoc test (P<0.05).

#### 3.3.2. Endophyte presence estimation in planta

At 75 dpi, P5 presence near the inoculation point was 4.81 times higher than in non-inoculated trees (*P*<0.05). At 50 cm, however, no significant differences were observed (*P*>0.05). The estimated presence of YM11 in inoculated trees was 9.09 times higher than in non-inoculated trees at 50 cm from the inoculation point (*P*<0.05) (Fig. 5A). YCB36 showed increased presence at both 5 and 50 cm above the inoculation point (4.64 and 17.86 times) (*P*<0.05) (Fig. 5A). In all cases, amplification of ITS sequences in non-endophyte inoculated trees occurred in later cycles for YCB36 than for YM11 and P5, indicating a higher basal presence of P5 and YM11 than YCB36 in non-endophyte-inoculated elms.

**Fig. 5:**
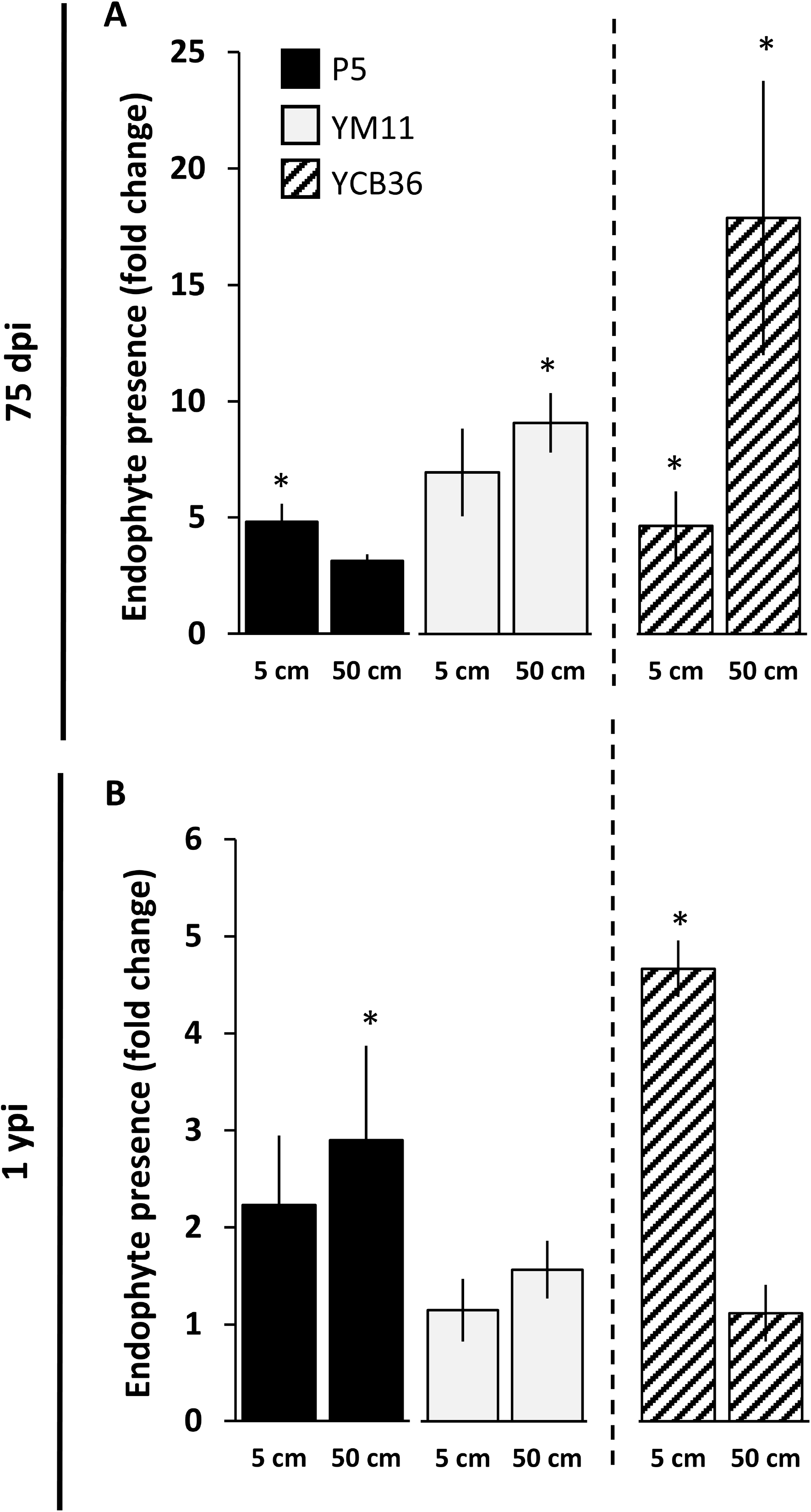
Relative presence of fungal endophytes estimated by qPCR at 75 dpi **(A)** and 1 ypi **(B)** in P5+YM11-inoculated trees (left panel) and YCB36 inoculated trees (right panel). Quantifications were performed at 5 and 50 cm above the endophyte inoculation point. Results are shown as a fold change of presence in non-endophyte inoculated trees (E-O-). Asterisks indicate significant differences to E-O- trees according to Fisher’s post-hoc test (P<0.05).

One year after inoculation, endophyte presence estimation was typically lower than in 75 dpi (Fig. 5B). P5 and YCB36 presence was still higher than in control plants at 50 and 5 cm from the inoculation point, respectively. Conversely, the presence of YM11 decreased to basal levels, showing no statistically significant differences with non-inoculated trees (*P* >0.05).

#### 3.3.3. Biochemical changes induced in endophyte-inoculated trees

Potential mechanisms of induced resistance were explored in YCB36-inoculated plants, as this was the only treatment that showed a protective effect against *O. novo-ulmi* in the trees. Soluble sugars and starch content *in planta* were not modified by endophyte inoculation (*P*>0.15). However, both flavonoids and total phenols were significantly modified by YCB36 treatment. Flavonoid content was higher in YCB36-inoculated trees than in controls at 50 cm from the inoculation point (*P*<0.05). Total phenols increased at both 5 and 50 cm from the inoculation point (*P*<0.05) (Fig. 6A, B). Proline content significantly decreased in endophyte-inoculated trees at 50 cm above the inoculation point (Fig. 6C).

**Fig. 6:**
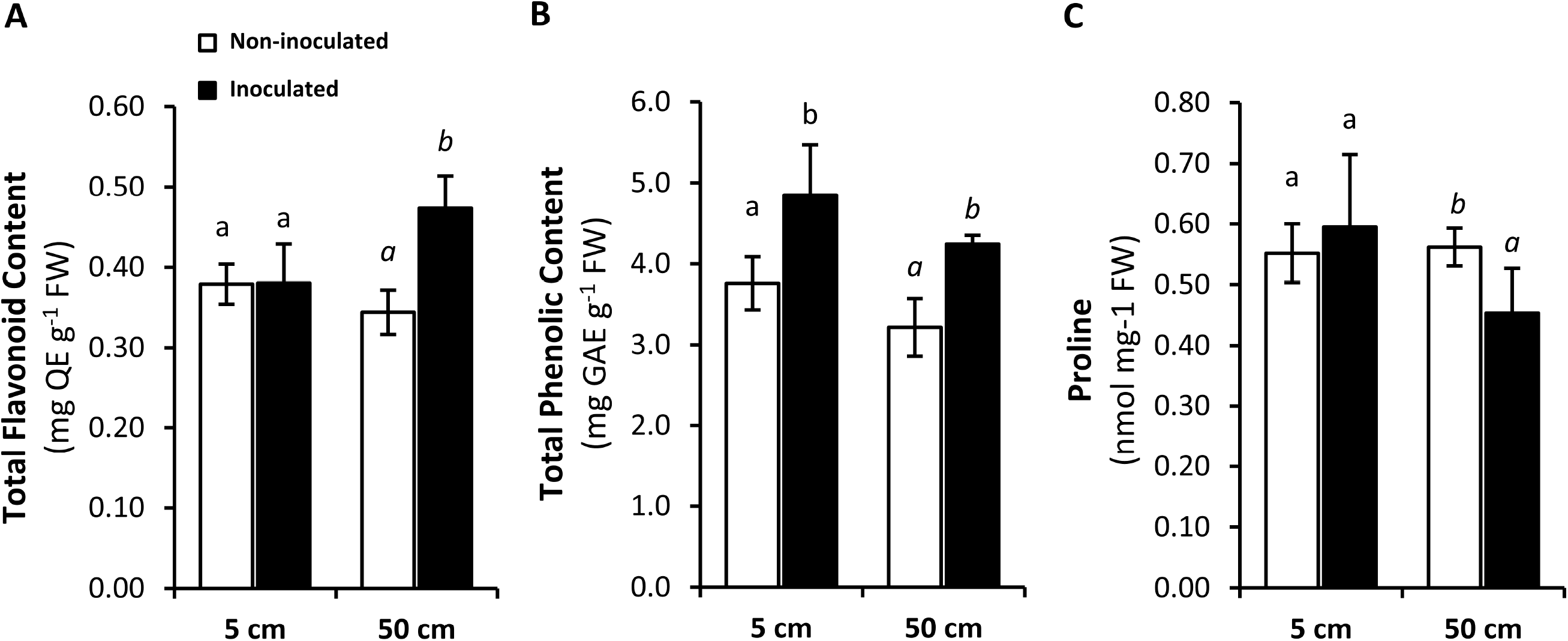
Biochemical measurements in wood samples of YCB36-inoculated and control trees at 5 and 50 cm from the inoculation point and at 75 dpi. **(A)** Total flavonoid content; (**B)** total phenolic content and **(C)** proline content. Different letters in the graphs indicate significant differences between YCB36-inoculated and non-inoculated trees at 5 cm (normal letters) and 50 cm (italics) according to Fisher’s post-hoc test (P<0.05).

## 4. Discussion

### 4.1. Phenotypic traits of the endophytic yeasts P5 and YM11

According to the phenotypic traits of the endophytes, P5 (*Cystobasidium*) can be classified within the group of plant growth-promoting yeasts (Joubert and Doty, 2018) and YM11 (*Exophiala*) within the dark septate endophytes (DSE) (Andrade-Linares and Franken, 2013). Endophytic yeasts have been shown to stimulate plant growth, increase yield and reduce biotic or abiotic damage (Punja and Utkhede, 2003; Kandel et al., 2017; de Tenorio et al., 2019). Plant experiments performed with the pink yeast *Rhodotorula* spp. demonstrated increased phytoremediation ability (Silambarasan et al., 2019), strong inhibition of plant fungal or bacterial pathogens (Kalogiannis et al., 2006; Akhtyamova and Sattarova, 2013), increased plant germination rates (Akhtyamova and Sattarova, 2013) and increased root and shoot growth rates, mainly through their ability to synthesize phytohormones (Nutaratat et al., 2014; Sen et al., 2019). On the other hand, DSE comprise a polyphyletic group of non-mycorrhizal ascomycetous fungi with potentially beneficial functions for their host plants, such as reducing biotic and abiotic stresses (Jumpponen and Trappe, 1998; Andrade-Linares, 2011; Tellenbach et al., 2013) and improving plant performance by increasing N and P acquisition (He et al., 2019) or synthesizing or modifying phytohormone levels *in planta* (Schulz and Boyle, 2005; Wu et al., 2020). Plant studies on *Exophiala* spp. revealed their ability to increase host tolerance to abiotic stress by activating antioxidant enzymes (Khan et al., 2012) and improving plant growth through production of biologically active GAs (Khan et al., 2009). In our work, however, the *Exophiala* isolate (YM11) did not produce significant amounts of phytohormones and was the endophyte with the lowest GA_3_ production, while P5 showed high production of IAA in the *in vitro* experiment (Fig. 3A). These results raise doubts about the role of YM11 as a producer of plant growth hormones, but support the classification of P5 as an IAA-producing yeast (Xin et al., 2009; Joubert and Doty, 2018). GA_3_ and IAA are the main hormones inducing cell proliferation and elongation, leading to enhanced root growth and stem elongation. Maintenance of high growth rates during *O. novo-ulmi* infection has been reported in small saplings as a resistance mechanism against the pathogen (Martín et al., 2019), and this mechanism could be enhanced in plants extensively colonized by hormone-producing yeasts. Although in the field assay the P5+YM11 treatment did not enhance tree resistance to DED, preliminary results with a similar P5+YM11 inoculum suggest that this treatment can induce root elongation (Martínez-Arias et al., unpublished results). Interestingly, YM11 and YCB36 were more inhibited than P5 by secondary metabolites that the tree might produce (Fig. 2), such as quercitin, cathechin, or gallic acid (Heimler et al. 1994; Martín et al., 2013). This suggests that P5 could be better adapted to grow under the chemical profile of elm wood.

Although both yeasts showed a weak inhibition of *O. novo-ulmi*, YM11 liquid filtrates were able to significantly reduce pathogen growth in liquid media, suggesting that inhibitory metabolites were secreted at a higher concentration in liquid culture than in solid medium.

### 4.2. Inhibition mechanisms of O. novo-ulmi by endophyte YCB36

YCB36 (Phaeosphaeriaceae) demonstrated strong *in vitro* antibiotic activity against *O. novo-ulmi.* The identity of this fungus at the species level using the ITS sequence is uncertain, but the most similar hits were found to be associated endophytically with a *Quercus* species (Moricca et al., 2012; Doust et al., 2017) (see NCBI accession numbers, Table 2). According to the LSU sequence, however, this strain can be tentatively assigned to *Hydeomyces* (Table 2). The activity of YCB36 against *O. novo-ulmi* in the *in vitro* dual culture assay evidenced strong pathogen inhibition without physical contact between mycelia, i.e. by antibiosis (Table 3; Fig. S2A, D). YCB36 liquid exudates (evaluated in the antibiosis assay) exerted the highest inhibition of *O. novo-ulmi*, supporting the previous results and suggesting release of antibiotic metabolites into the culture medium (Table 3; Fig. S2C). The high hydrolytic activity of this endophyte may be involved in *O. novo-ulmi* growth inhibition. Both chitinolytic and proteolytic activities were highly positive in YCB36 (Table 3; Fig. S3), indicating that this fungus may have released these enzymes into the culture medium. Chitin is the major structural component of hyphal cell walls, and its enzymatic degradation via chitinases can disrupt or collapse fungal hyphae or even cause fungal death (Huang et al., 2007). For this reason, chitinolytic microorganisms have a potential application as biological control agents, as is the case with *Trichoderma sp.* (Aoki et al., 2020). In turn, proteases are usually needed to facilitate fungal entry into host tissues, and also play an important role in pathogen virulence during the entire infection cycle (Hueck, 1998; Figaj et al., 2019). In this experiment, as expected, the highest proteolytic activity was observed in *O. novo-ulmi*. This feature may give the pathogen an advantage over other fungi by enabling it to rapidly colonize elm xylem tissue. YCB36 also showed positive protease activity that could facilitate plant colonization and also affect *O. novo-ulmi* growth (Zhang et al., 2012).

Emission of antimicrobial VOCs by YCB36 may be another reason for the reduction in *O. novo-ulmi* growth. Several endophytic fungi have been reported to emit VOCs to establish a communicative interaction with their surroundings (Farre-Armengol et al., 2016; Roy and Banerjee, 2019). These molecules could have a potential role against fungal diseases due to their antimicrobial effects (Ezra et al., 2004; Chen et al., 2018; Wonglom et al., 2020) and/or ability to induce expression of defence genes in plants (Malmierca et al., 2012; Naznin et al., 2014; Pieterse et al., 2014). YCB36 showed the ability to reduce *O. novo-ulmi* growth when grown in confronted plates, probably through emission of volatile antibiotic molecules commonly attributed to terpenes, alcohols, and carboxylic acids (Zhang et al., 2014). Both chitinase and VOC emission by YCB36 could have a negative influence on *O. novo-ulmi* development in plant tissues and may have contributed to the reduced DED symptoms observed in YCB36-inoculated trees. Furthermore, a high nutritional niche overlap was observed between *O. novo-ulmi* and YCB36 in the nutrient utilization assay. Contrary to the results obtained by Blumenstein et al. (2015) with other elm endophytes, *O. novo-ulmi* and YCB36 showed the same range of metabolized carbon substrates, indicating a high nutritional niche overlap that could result in competitive interaction between endophyte and pathogen *in planta*. However, unlike YCB36, *O. novo-ulmi* is able to grow in the presence of certain inhibitory substances such as gallic acid and thymol, which could reflect its greater ability to cope with plant defensive metabolites.

Additionally, both *O. novo-ulmi* and YCB36 (in particular) displayed the ability to produce siderophores. Microorganisms use siderophores to obtain iron from the environment. In pathogenic interactions, they enable iron acquisition from host proteins and have an essential role in virulence in several plant pathogens (Oide et al., 2006; Chen et al., 2013; Albarouki et al., 2014). In mutualistic symbiotic relationships between plants and fungi, siderophores produced by the fungal symbiont facilitate plant access to natural insoluble iron found in soil, improving plant growth (Grobelak and Hiller, 2017). Many studies have also described the role of siderophores as stimulators of plant immunity (Buysens et al., 1996), in particular as activators of induced systemic resistance (ISR) (Beneduzi et al., 2012; Aznar and Dellagi, 2015), because they can be recognized as microbe-associated molecular patterns (MAMPs). Microbial gene-knockout on siderophore emission revealed the importance of these molecules for ISR elicitation (De Vleesschauwer et al., 2008).

### 4.3. Endophyte establishment and persistence in host tissues

In the *in vivo* experiment, endophyte inoculations were performed for the first time on elm trees using the BITE® endotherapy device. Fungal DNA amplification on E+O- wood samples confirmed the success of the inoculation system, with inoculated trees showing higher P5, YM11 or YCB36 fungal presence than in non-inoculated trees (Fig. 5A, B). Nevertheless, the increased presence of these endophytes led to reduced symptoms after *O. novo-ulmi* inoculation only in YCB36-inoculated trees (Fig. 3C). YCB36 showed a higher plant colonization rate than P5 and YM11 (Fig. 5A, B), possibly enhanced by its high protease activity. In the long term (one year after inoculation), however, the presence of YCB36 in plant tissue was markedly reduced, and the YCB36 isolate no longer exerted a protective effect (Fig. 5D), similar to what occurs in preventive inoculations with *Verticillium albo-atrum* in *Ulmus* sp. (Postma and Goossen-van de Geijn, 2016). In our case, despite the use of local elm endophytes, the elm fungal endobiome was resilient enough to reduce the presence of the artificially inoculated endophytes. This corroborates the assumption that the microbiome tends to be resistant and resilient to disturbances in community structure and has a composition that is very difficult to alter (Toju et al., 2018). The strength of microbiome communities seems to arise from pioneering fungi colonizing the seed or the plant at early plant developmental stages, when they are able to use space and resources and limit or slow down the establishment of later colonizers (Fukami et al., 2015, Wei et al., 2015;). Therefore, managing the microbiome at early developmental stages, i.e. before or during seed germination, using the targeted endophytes as early colonizers may be a worthwhile strategy for future experiments.

### 4.4. Effects of YCB36 inoculation on tree physiology and defence metabolism

Interestingly, YCB36-inoculated trees (E+O-) showed increased leaf stomatal conductance (Fig. 4B), suggesting that this endophyte has the ability to modulate plant physiological functions. Several studies have highlighted the considerable influence of microbial symbionts on host water relations (Rho and Kim, 2017, and references therein). Stomatal conductance is positively related to photosynthetic CO_2_ uptake and, consequently, primary metabolism, which is involved in plant fitness and defence capability (Rosa et al., 2009). In this assay, no significant modifications in non-structural carbohydrates were observed, although we noticed a higher level of secondary metabolites (flavonoids and total phenolic content) in YCB36-inoculated trees than in non-inoculated trees (Fig. 6C, D). Phenolic compounds are widely distributed in plants and play key roles in metabolic and physiological processes (Boudet, 2007). These molecules also have antioxidant properties (Huang et al., 2007) that can ameliorate plant performance under stress conditions (Daayf et al., 2012; Cheynier et al., 2013; Mishra et al., 2018). Several studies have demonstrated increased synthesis of flavonoids and phenols in plants colonized by symbiotic microorganisms (Qawasmeh et al., 2012; Dupont et al., 2015). White et al. (2010) defended the hypothesis that enhanced production of antioxidants in endophyte-inoculated plants may be due to the plant response to endophyte-produced reactive oxygen species (ROS). Increased phenolic metabolites can also contribute to enhanced tree resistance to *O. novo-ulmi* in YCB36-inoculated trees, as reported in other elm studies (Witzell and Martín, 2008; Li et al., 2016). Moreover, accumulation of phenolic compounds may be linked to ISR activation. As described earlier, elm defence response could be triggered by YCB36 MAMPs, such as siderophores and VOCs. ISR activation is often regulated by a jasmonic/ethylene-dependent signalling pathway, and culminates with synthesis of pathogenesis-related proteins (PR proteins), synthesis of phytoalexins or, as observed in the YCB36-inoculated plants, accumulation of phenolic compounds (Conrath at al., 2002). Proline content reduction in YCB36-inoculated trees may also indicate reduced tree stress. Lower proline levels have been described in various beneficial plant-microbe interactions, sometimes associated with better nutritional status and fitness and lower stress levels (Verslues and Sharma, 2010; Martínez-Arias et al., 2020).

Three different mechanisms may have contributed to the reduction in DED symptomatology in YCB36-inoculated trees: i) induction of plant resistance mechanisms that indirectly inhibit *O. novo-ulmi* development (ISR activation and phenolic accumulation); ii) *O. novo-ulmi* niche displacement by YCB36 due to nutritional overlap; and iii) direct inhibition of *O. novo-ulmi* by YCB36 through chitinase, antifungal metabolites, and VOC emissions. These mechanisms could act independently or synergistically. Examples include the symbiotic bacteria *Pseudomonas aeruginosa*, which is able to trigger ISR either through siderophore (Audenaert et al., 2002) or volatile emission (Ryu et al., 2004), and the *Trichoderma* biocontrol fungi, which have been shown to activate defence genes through VOC emission and inhibit pathogen growth by interacting with hydrolytic enzymes (Malmierca et al., 2012).

## Conclusions

Our study provides new evidence for the role of fungal endophytes as alleviators of pathogen damages and for the mechanisms underlying this activity. YCB36, the endophytic fungus with highest antibiotic, protease, chitinase and siderophore activities in *in vitro* tests, significantly reduced DED symptomatology in the field test. Therefore, this endophyte is expected to be a promising candidate for development into a biological control agent to prevent or reduce DED damages. In contrast, although both endophytic yeasts were successfully established in elm stems after artificial inoculation into adult trees, neither *Cystobasidium* sp. (P5) nor *Exophiala* sp. (YM11) isolates alleviated wilting symptoms,

Further studies are needed to improve future field applications of YCB36 in elm trees. Because we observed a reduced YCB36 load one year after its application, annual or more frequent applications of the target endophyte could be necessary. Inoculation of YCB36 into elm seeds or in early developmental stages of vegetative reproductive materials (e.g., *in vitro*-propagated plants) could be another interesting strategy for producing plants colonized by the beneficial endophyte, since early plant colonizers have the advantage of using the space and resources earlier than subsequent competitors (priority effects). Moreover, similarly to the strategy followed by Miller et al. (2008), Tellenbach et al. (2013) or McMullin et al. (2018) for *Phialocephala* sp. endophytes, further information about the antibiotic molecules released by YCB36 *in vitro* is required to understand a mechanistic basis of YCB36 activity against DED. Additionally, more efficient protocols, including different inoculum densities and other inoculation methods, need to be examined to optimize treatments. Other elm clones and environmental conditions should be tested to validate the results, because previous research demonstrated the significance of these variables in the stability of tree protection against DED that some endophytes confer (Martín et al., 2015). Also, high-throughput methods for conidia production need to be examined to obtain mass quantities of cells, as conidia are the most resilient and suitable cell type used in formulations for biocontrol treatments (Mancera-López et al., 2019). Future experiments must be specifically planned to plan future large-scale application strategies.

## Acknowledgments

We thank Jorge Dominguez, David Medel and David Macaya for assistance in plant propagation and endophyte isolation. This study was funded by the project GENESIS (AGL-2015-66952-R; MINECO/ERDF) and by an agreement for elm conservation between Universidad Politécnica de Madrid and *Dirección General de Desarrollo Rural y Política Forestal* (MAPA/EAFRD). C.M-A was supported by an FPI pre-doctoral fellowship from the Spanish Ministry of Economy and Competitiveness.

---

**Fig. S1:**
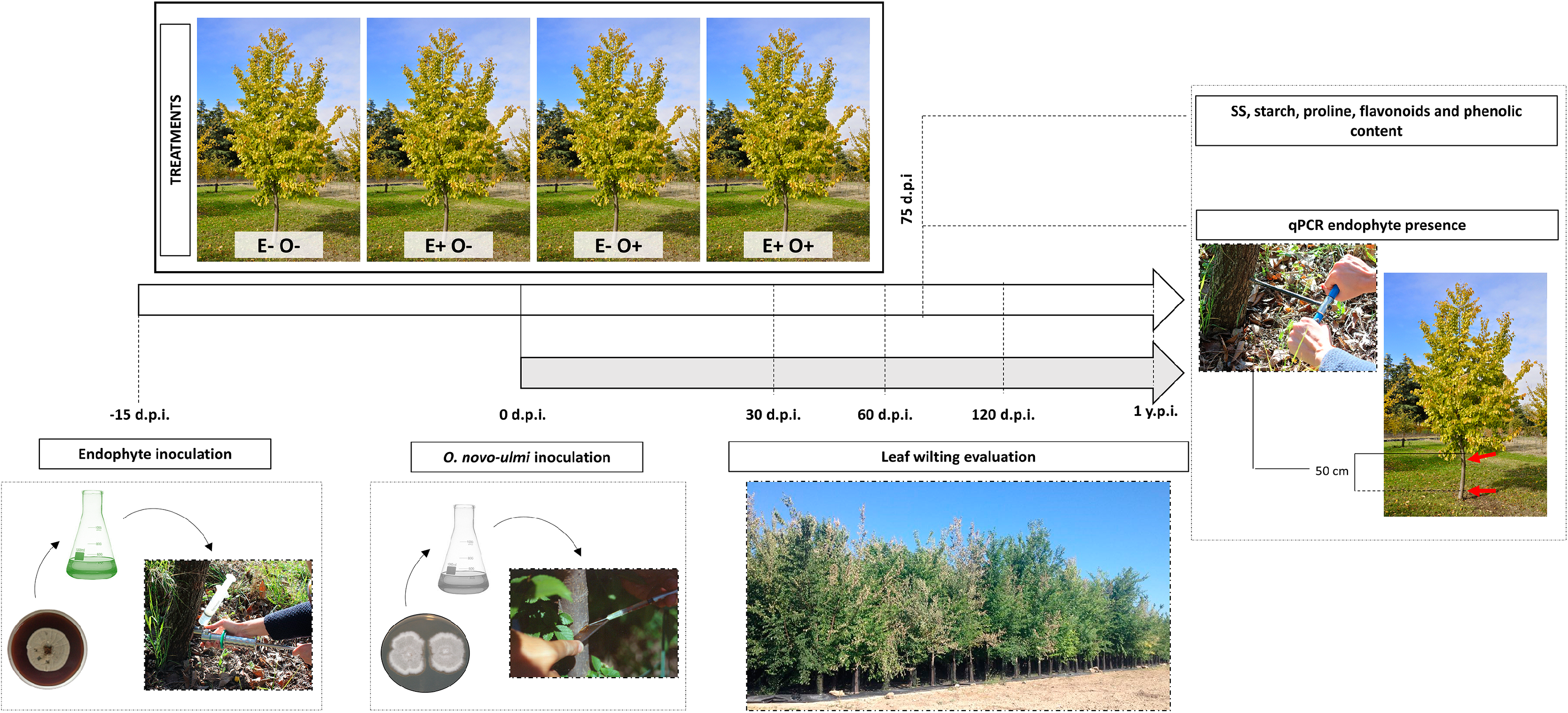
Experimental procedure. First, a fungal cell suspension was inoculated into the vascular system at the trunk base using an endotherapy device. *O. novo-ulmi* inoculations were performed 15 days later using a spore suspension following traditional inoculation methods for DED. Four treatments were applied for each endophyte combination (P5+YM11 or YCB36): E+O- : endophyte-inoculated plants without *O. novo-ulmi* inoculation, E-O+: *O. novo-ulmi*-inoculated plants without previous endophyte inoculation, E+ O+: endophyte and *O. novo-ulmi-*inoculated plants, and E-O-: non-inoculated plants (neither endophyte nor *O. novo-ulmi*). Leaf wilting was evaluated in each tree at 30, 60, 120 days post (pathogen) inoculation (dpi) and 1 year post-inoculation (ypi) by three independent assessors. At 75 dpi a wood sample was cored with an increment borer (N=3) at 5 and 50 cm above the inoculation point for biochemical and fungal quantification measures. SS = soluble sugars.

**Fig. S2:**
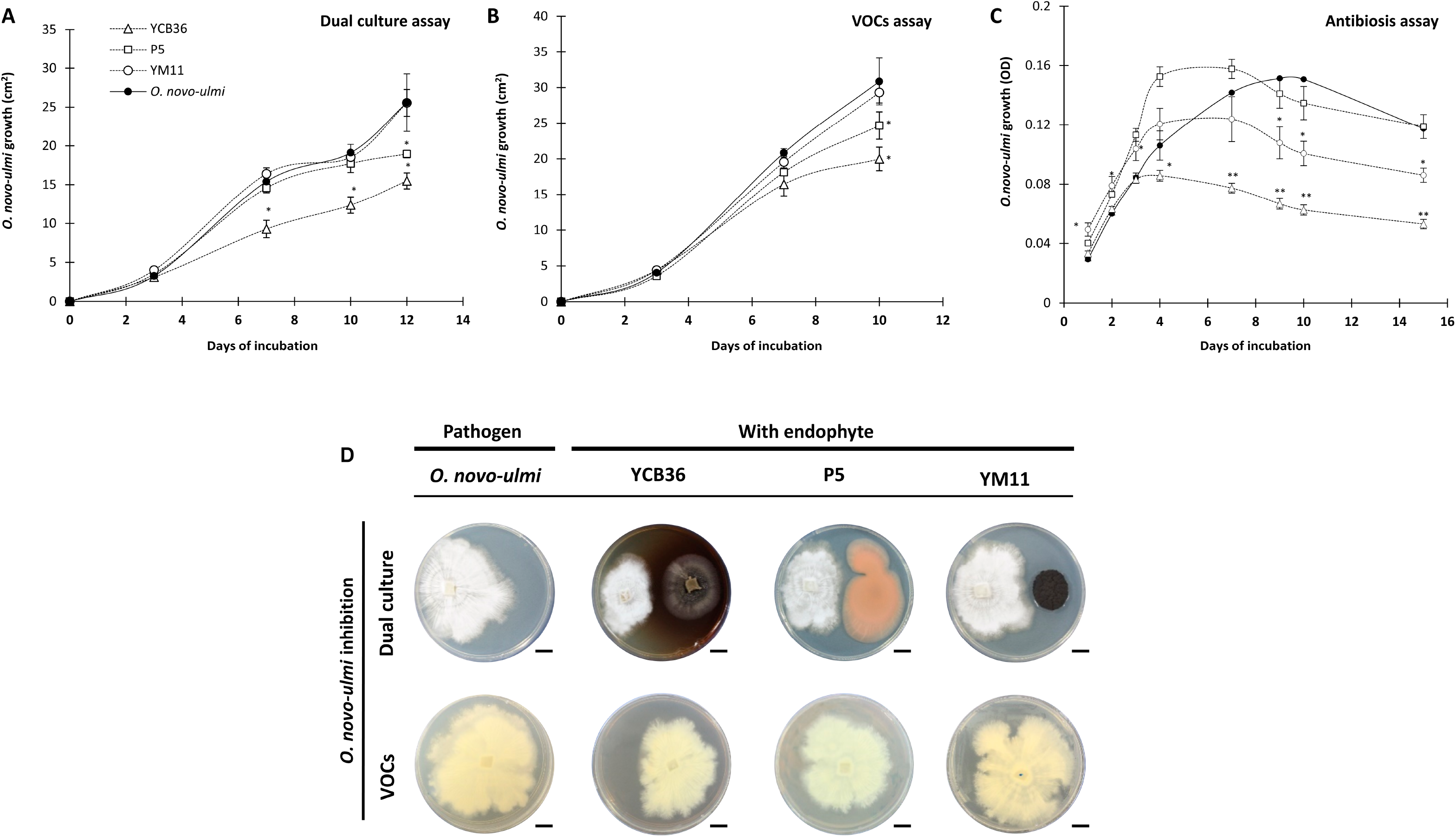
*In vitro* interactions between *O. novo-ulmi* and fungal endophytes. (A) *O. novo-ulmi* growth when confronted with fungal endophytes in dual late assays; (B) *O. novo-ulmi* growth when confronted with fungal endophytes in petri dishes to evaluate the effect of endophyte volatile substances (VOCs) emission; (C) *O. novo-ulmi* growth in the presence of endophyte liquid filtrates in 96-well microplates. In (C) *O. novo-ulmi* growth is measured by optical density (OD). Asterisks indicate statistically significant differences to the control treatment according to Fisher’s post-hoc test (P<0.05). Dual culture and VOC assay illustrations (D). In the dual culture, *O. novo-ulmi* (white colony at the left side of the plate) is confronted with fungal endophytes (YCB36, P5 and YM11) (right side of the plate) at day 12. The VOCs assay shows the results of *O. novo-ulmi* growth in the presence of endophytes in confronted plates at day 10. The scale bar below each petri dish indicates a length of 1 cm.

**Fig. S3:**
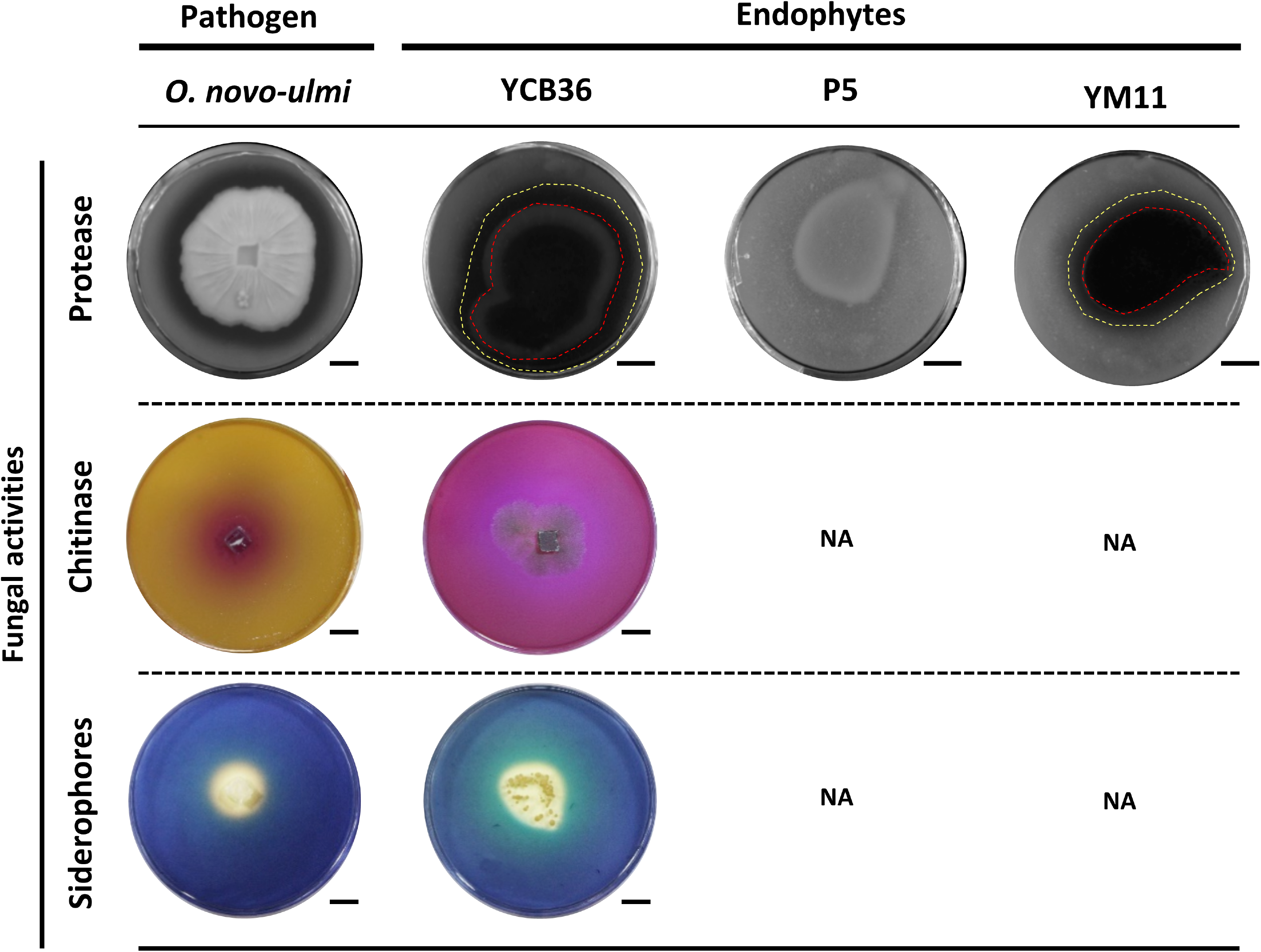
Protease and chitinase activities and siderophore production in *in vitro* assays performed by fungal endophytes and pathogen at day 20. Dotted red line delimits colony diameter; dotted yellow line delimits clear halo diameter. Chitinase activity and siderophore production were negative (NA) for endophytes P5 and YM11. The scale bar below each petri dish indicates a length of 1 cm.

